# Identification of genes required for enzalutamide resistance in castration-resistant prostate cancer cells *in vitro*

**DOI:** 10.1101/2020.03.27.011825

**Authors:** Sarah E. Kohrt, Wisam N. Awadallah, Robert A. Phillips, Thomas C. Case, Renjie Jin, Jagpreet S. Nanda, Xiuping Yu, Peter E. Clark, Yajun Yi, Robert J. Matusik, Philip D. Anderson, Magdalena M. Grabowska

## Abstract

Castration-resistant prostate cancer can be treated with the anti-androgen enzalutamide, but responses and duration of response are variable. To identify genes that support enzalutamide resistance, we performed a short hairpin RNA (shRNA) screen in the bone-homing, castration-resistant prostate cancer cell line, C4-2B. We identified eleven genes (*TFAP2C, CAD, SPDEF, EIF6, GABRG2, CDC37, PSMD12, COL5A2, AR, MAP3K11*, and *ACAT1*), whose loss resulted in decreased cell survival in response to enzalutamide. To validate our screen, we performed transient knockdowns in C4-2B and 22Rv1 cells and evaluated cell survival in response to enzalutamide. Through these studies, we validated three genes (*ACAT1, MAP3K11*, and *PSMD12*) as supporters of enzalutamide resistance *in vitro*. Although *ACAT1* expression is lower in metastatic castration-resistant prostate cancer samples versus primary prostate cancer samples, knockdown of *ACAT1* was sufficient to reduce cell survival in C4-2B and 22Rv1 cells. *MAP3K11* expression increases with Gleason grade, and the highest expression is observed in metastatic castration-resistant disease. Knockdown of *MAP3K11* reduced cell survival and pharmacologic inhibition of MAP3K11 with CEP-1347 in combination with enzalutamide resulted in a dramatic increase in cell death. This was associated with decreased phosphorylation of AR-Serine650, which is required for maximal AR activation. Finally, while *PSMD12* expression did not change during disease progression, knockdown of *PSMD12* resulted in decreased AR and AR splice variant expression, likely contributing to the C4-2B and 22Rv1 decrease in cell survival. Our study has therefore identified at least three new supporters of enzalutamide resistance in castration-resistant prostate cancer cells *in vitro*.

**Financial support:** The authors would like to acknowledge funding from the Joe C. Davis Foundation (to RJM), the Vanderbilt Institute for Clinical and Translational Research (VICTR, to YY, PEC, and RJM). The Vanderbilt Institute for Clinical and Translational Research (VICTR) is funded by the National Center for Advancing Translational Sciences (NCATS) Clinical Translational Science Award (CTSA) Program, Award Number 5UL1TR002243. The content of this manuscript solely the responsibility of the authors and does not necessarily represent the official views of the NIH. We would also like to acknowledge the Case Research Institute, a joint venture between University Hospitals and Case Western Reserve University, start-up funds (to MMG), and the Cell and Molecular Biology Training Program (T32 GM 008056 to SEK).

## Introduction

Early stage prostate cancer is an androgen-dependent disease, and advanced prostate cancer is largely treated with androgen deprivation therapy, which targets androgen receptor (AR). Inevitably, tumors progress to castration-resistant prostate cancer. In castration-resistant prostate cancer, AR signaling continues through multiple mechanisms including AR full length (AR-FL) over-expression, increases in androgen production, and induction of constitutively active AR splice variants (AR-V)[1]. Currently, abiraterone acetate and enzalutamide are standard therapies in metastatic castration-resistant prostate cancer[2, 3]. Enzalutamide is well tolerated and effective, extending castration-resistant prostate cancer patient survival by 5 months versus placebo after chemotherapy[4] and delaying initiation of chemotherapy by 18 months versus placebo[5]. Nevertheless, tumors progress to enzalutamide-resistant castration-resistant prostate cancer. How these tumors progress to resistance is an area of active investigation, but both AR-dependent and AR-independent mechanisms have been proposed.

The purpose of our studies was to identify genes that support resistance to enzalutamide and identify gene products (proteins) that once targeted could re-sensitize tumors to enzalutamide. We first selected a bone metastasis-derived castration-resistant prostate cancer model, C4-2B[6], which upregulates AR-FL and AR-V7 in response to continuous enzalutamide treatment[7]. We then performed a short hairpin RNA (shRNA) screen, comparing the response of vehicle-treated and enzalutamide-treated C4-2B cells and assessed which genes, when knocked down, resulted in increased cell death. Through this approach, we identified 11 genes (*TFAP2C, CAD, SPDEF, EIF6, GABRG2, CDC37, PSMD12, COL5A2, AR, MAP3K11*, and *ACAT1*), of which *ACAT1, MAP3K11*, and *PSMD12* appear the most promising in validation studies.

## Methods

### shRNA screen

The 27k Module 1 DECIPHER lentiviral shRNA library (Cellecta, DHPAC-M1-P) was transduced into C4-2B cells, targeting approximately 5,043 genes, whose gene products represent members of signal transduction pathways and approved drug targets. Cells were transduced with a pooled lentiviral shRNA library, generated per manufacturer’s instructions, and then split into three groups. Cells in the first group were collected and represent the initial population of shRNA quantity (3095 YY1-2). The cells from the second group served as an enzalutamide-negative control group (cell culture in the enzalutamide vehicle control [DMSO] for 6 days; YY3095 3-5), and cells from the third group were cultured in the presence of 100nM enzalutamide for 6 days (3095 YY6-8). Genomic DNA was isolated and shRNA was quantified by deep sequencing. The ratio of the abundance of each shRNA in the third group versus both control groups was calculated. A given shRNA was considered a ‘‘hit’’ if it showed at least a 2-fold abundance decrease relative to both enzalutamide-negative control and initial samples.

### Computational workflow

We performed quality control using on the sequencing data using FastQC[8]. Using the DECIPHER Data and Pathway Analysis Software http://www.decipherproject.net/software/ called Bar-code Analyzer and Deconvoluter, web converted raw high-throughput sequencing data into a summary file for subsequent processing. This file includes annotation for every identified gene in our screen. For most transcripts being silenced, there are five barcodes from two different shRNA molecules targeting the transcript. Next, we used EdgeR [9, 10] and Limma[11, 12] to generate differential expression based on gene (barcode) count. After verifying that all of our samples had a similar sequencing depth, we used the Limma-trend test to identify which barcodes are decreased in enzalutamide-treated versus vehicle treated and untreated cells. Additional information can be found in the bioinformatics supplement (Supplemental Methods).

### Human clinical data

To assess the expression of our screen-identified genes during prostate cancer progression, we used the Prostate Cancer Transcription Atlas web tool (http://www.thepcta.org/), that includes 1,321 clinical specimens from 38 prostate cancer cohorts, detailed here[13]. Publication-ready images were downloaded. To evaluate the frequency of alterations of our putative drivers in metastatic castration-resistant prostate cancer, we used cBioPortal[14, 15] to interrogate a set of publically available data [16] with associated outcome and gene expression data [17]. This data set also includes an AR activity score, which is a measure of how active AR is, as well as a neuroendocrine prostate cancer (NEPCa) score, a measure of how neuroendocrine-like the gene expression pattern is. For the i) gene expression comparison between naïve and exposed patients and ii) correlation with AR and NEPCa activity scores, raw data was downloaded and plotted using GraphPad Prism. All other plots were exported as is from cBioPortal.

### siRNA transient transfections

22Rv1 cells were purchased from American Type Culture Collection (ATCC, CRL-2505). C4-2B[6] cells were provided by Drs. Ruoxiang Wu and Leland Chung (Cedars-Sinai) and Short Tandem Repeat (STR) validated. Both cell lines were mycoplasma tested following expansion of stocks. C4-2B and 22Rv1 cells were cultured in RPMI 1640 with L-glutamine in 10% fetal bovine serum. Cells were transfected with two siRNAs (Ambion Silencer Select, Thermo Fisher, see Table 1) per gene in Lipofectamine RNAi Max transfection reagent (Invitrogen) and OptiMEM medium (Gibco, Life Technologies). For cell survival assays, cells were transfected with 1 pmol siRNA for 24 hours then cells were either treated with 1:1000 DMSO (vehicle) or 10µM enzalutamide for six days in RPMI medium, with a media change at day three. For protein expression studies, cells were transfected with 30 pmol siRNA for 3 days or 6 days.

**Table 1:** siRNA constructs.

| **Name** | **Cat No. #1** | **Cat No. #2** |
| --- | --- | --- |
| siNT | 4390843 | 4390846 |
| **Name** | **Gene ID #1** | **Gene ID #2** |
| siAR | s1539 | s1538 |
| siACAT1 | s900 | s901 |
| siMAP3K11 | s8816 | s8814 |
| siGABRG2 | s532146 | s532148 |
| siPSMD12 | s11417 | s11418 |
| siCDC37 | s21967 | s21965 |
| siCOL5A2 | s3310 | s533841 |
| siSPDEF | s195114 | s24518 |
| siCAD | s2139 | s224811 |
| siEIF6 | s7586 | s7587 |
| siTFAP2C | s14011 | s14009 |

### Quantitative PCR (qPCR)

Total RNA was extracted from cells using the RNeasy Plus Mini kit (Qiagen) according to manufacturer instructions. RNA was reverse transcribed to cDNA using Bio-Rad iScript cDNA Synthesis Kit following manufacturer instructions. Following reverse transcription, qPCR was performed to quantitate RNA levels in C4-2B and 22Rv1 cells and to establish efficacy of siRNA constructs. qPCR was performed using the SYBR™ GREEN PCR Master Mix kit (Bio-Rad). Relative expressions of total RNA were normalized to endogenous control Glyceraldehyde 3-phosphate dehydrogenase (GAPDH). All samples were run with two technical replicates and six biological replicates. No reverse transcriptase controls and no transcript controls were run in duplicate along with each primer in every experiment. Unpublished primers were designed using Primer-BLAST[18] to span exon-exon junctions when possible and identify as many transcript isoforms as possible. All primers were tested for amplification efficiency by using a standard curve generated from four 1:10 serial dilutions of cDNA prior to experiments. Efficiency was calculated using the formula E = −1+10(−1/slope) where E=Efficiency. All primers used reported >80% efficiency according to this method. Primers are listed in Table 2.

**Table 2:**
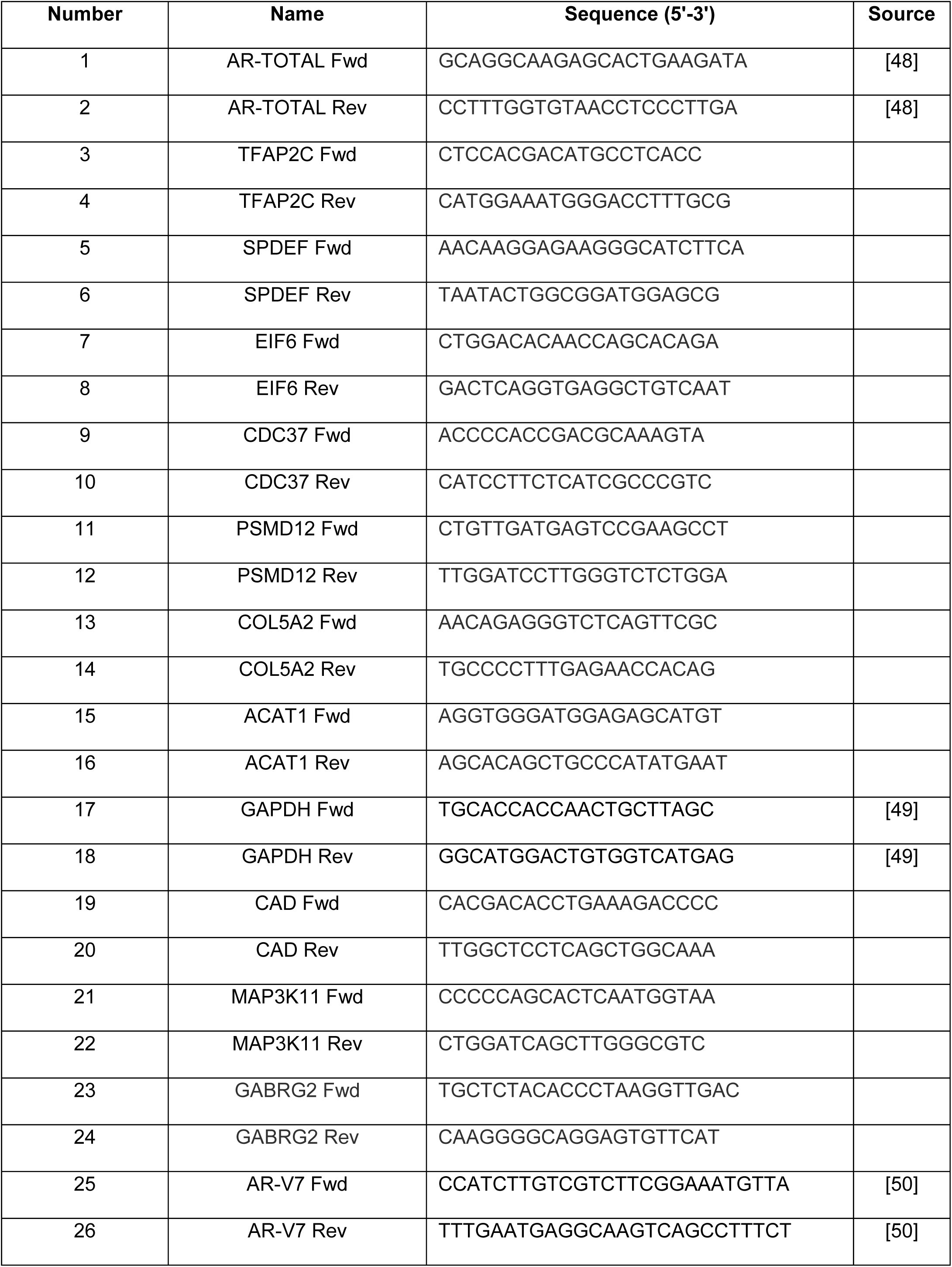
qPCR primers.

### Cell survival assays

Cell survival in response to knockdowns and enzalutamide was assessed using crystal violet staining. The crystal violet staining protocol was used as previously described [19]. In brief, 1×10^4^ cells were seeded per well in 100uL RPMI 1640 with L-glutamine in 10% fetal bovine serum and underwent described drug treatments with eight biological replicates. At the end of the treatment course, wells were fixed in 4% paraformaldehyde in PBS (Alfa Aesar) and stained with 0.05% crystal violet stain (Ricca). Cells were washed with deionized water and dried for 16-24 hours before being imaged. Destaining for quantification was performed using 10% acetic acid and absorbance was read at 590 nm using a spectrophotometer (SpectraMax iD3, Molecular Devices). An average value of non-targeting vehicle was generated, and each value was divided by this average value to generate a normalized fold-change cell survival reading.

### Western blotting

Protein isolation was performed using working RIPA buffer (120mM NaCl, 50 mM pH 8.0 Tris, 0.5% NP-40, 1 mM EGTA) containing protease and phosphatase inhibitors (100 ug/mL PMSF, 1 mM NaOrVa, 50 ug/mL aprotinin, 50ug/mL leupeptin). Protein concentration was determined using a Bio-Rad protein concentration assay dye and absorbance read at 595 nm on a spectrophotometer (SpectraMax ID3, Molecular Devices). 20µg total protein in βmercaptoethanol-containing loading buffer was added per well onto 4-12% SDS-PAGE gels (NuPAGE) and transferred to PVDF (Bio-Rad) or nitrocellulose (Amersham) membrane. Nitrocellulose membranes were stained for total protein using Ponceau S (Boston BioProducts). Membranes were blocked with 10% BSA or milk-fat in TBST for 1 hour at room temperature. Primary antibodies from Abcam for AR (ab74272), AR-V7 (ab198394), phosphorylated AR-Ser650 (ab47563), ACAT1 (ab168342), MAP3K11 (ab51068), PSDM12 (ab229930), and GAPDH (AM4300) were used at recommended concentrations in 2.5% milk-fat or 2.5% bovine serum albumin overnight at 4°C. Anti-rabbit and anti-mouse secondary antibodies (GE Healthcare) were used in 2.5% milk-fat for one hour at room temperature. Membranes were imaged via chemiluminescence reagents (Super Signal™ West Pico Plus, Thermo Fisher) on a digital imager (ChemiDoc™ Touch, Bio-Rad).

### Drug treatments

Cell response to the MAP3K11 inhibitor CEP-1347 was assessed in crystal violet cell survival assays. 1×10^4^ cells were seeded per well in 100uL RPMI 1640 with L-glutamine in 10% fetal bovine serum and allowed to settle. CEP-1347 (Tocris, Catalog no. 4924) reagent was resuspended in DMSO for 1mM stock. Working solutions of 80μM, 40μM, 20μM, and 10μM were made with DMSO, then diluted 1:100 in RPMI 1640 with L-glutamine in 10% fetal bovine serum for final concentrations of 800nM, 400nM, 200nM, and 100nM, respectively. Enzalutamide (MDV3100; Selleckchem CAS No. 915087-33-1) was resuspended in DMSO for a working concentration of 100mM then diluted 1:1000 in media. DMSO control media contained equal amounts of solvent as drug treatment samples. 100uL of drug-containing media was plated per well for six days with new drug treatments performed on the third day. Cell survival at the end of drug treatment course was assessed via crystal violet staining as described.

### Statistical analysis

Statistical analysis was performed using GraphPad Prism. For analysis of cell survival experiments, following the removal of outliers using the ROUT method, vehicle to vehicle, enzalutamide to enzalutamide comparisons were made using the Kruskal-Wallis test as data was not always normally distributed. Comparisons between gene expression of abiraterone and/or enzalutamide exposed and naïve patients used the non-parametric Kruskal-Wallis test as data was not normally distributed. Correlations between gene expression (FPKM capture) and AR and NEPCa activity scores were calculated using non-parametric Spearman correlations. Clinical data from the Prostate Cancer Transcription Atlas web tool included statistical analysis.

## Results

To identify genes that support enzalutamide resistance in castration-resistant prostate cancer *in vitro*, we transduced C4-2B cells with a lentiviral shRNA library that targeted 5,043 genes whose products act in signal transduction and are drug targets. The products of these genes include kinases, enzymes, receptors, and other proteins that already have targeted therapies. C4-2B cells were transduced with the shRNA library and split into three groups, one collected at day zero, one treated with vehicle for six days, and one treated with enzalutamide for six days. We used high-throughput sequencing to quantitate the number of shRNA barcodes in each sample. We were predominantly interested in the identity of the shRNA barcodes that had decreased in abundance in the enzalutamide-treated cells versus non-treated and vehicle-treated cells, as the loss of these barcodes reflected the death of cells in which these genes were essential. Through this analysis we identified 11 genes whose silencing resulted in increased cell death in response to enzalutamide: *TFAP2C, CAD, SPDEF, EIF6* (ITGB4BP), *GABRG2, CDC37, PSMD12, COL5A2, AR, MAP3K11*, and *ACAT1* (Figure 1A). As prostate cancer is driven by AR, and enzalutamide targets AR, the presence of *AR* in our gene list was an encouraging sign. The fold change differences were more dramatic between shRNA counts from enzalutamide-treated cells versus the initial population of shRNAs (Figure 1B) as compared to shRNA counts from vehicle treated cells (Figure 1C). This likely reflects the importance of some of these genes in cell survival in general and/or in response to DMSO.

**Figure 1:**
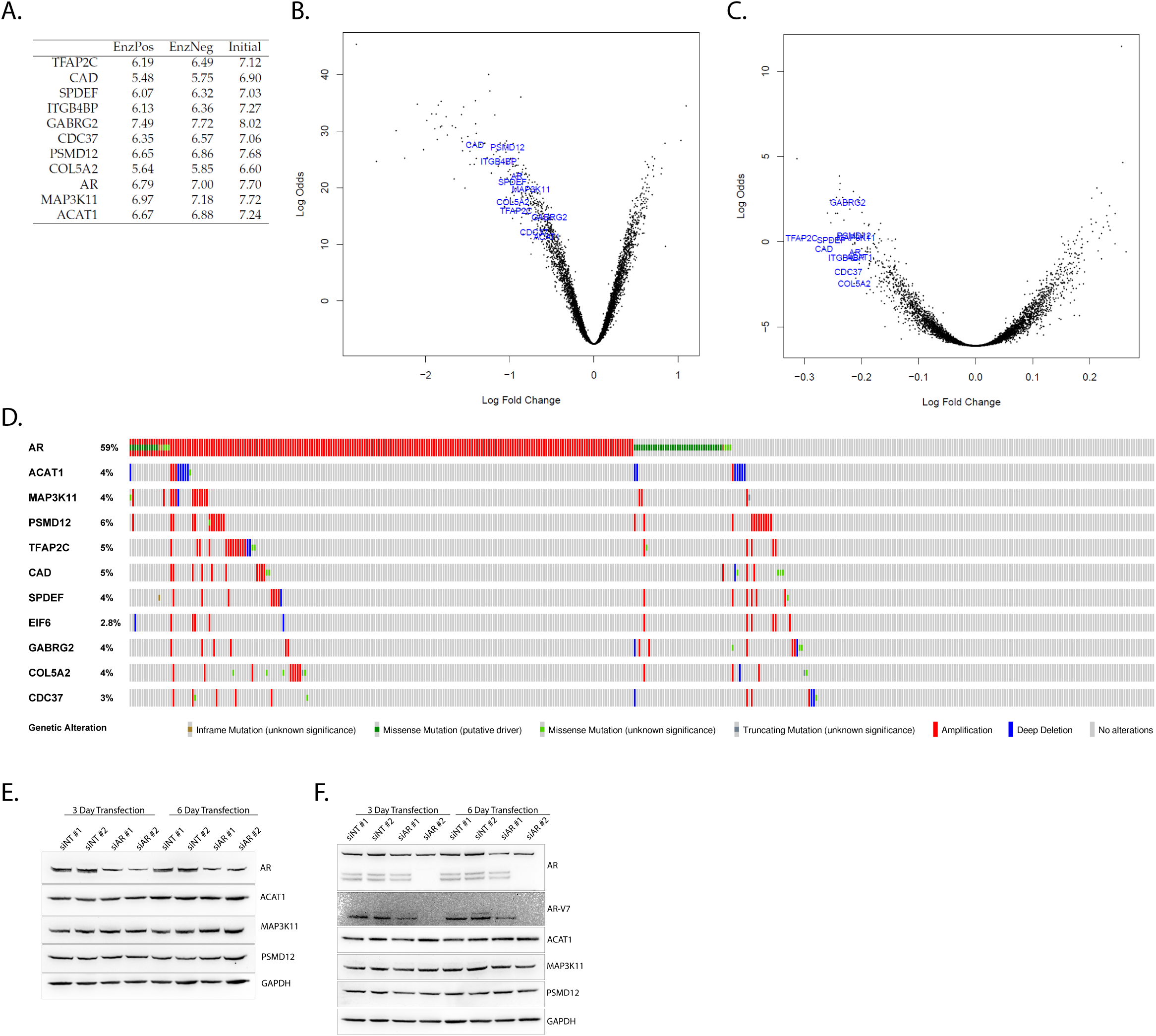
Identification of genes that support enzalutamide resistance in C4-2B cells. **A**. Log-scaled shRNA barcode counts of differentially expressed shRNA barcodes (genes) in C4-2B cells in response to enzalutamide (EnzPos) and vehicle (EnzNeg). Cells collected at the start of the experiment (Initial) represent the starting pool of shRNA barcodes. ITGB4BP is now EIF6. **B/C**. Under-represented shRNA barcodes (genes) between enzalutamide-treated and untreated (**B**) and vehicle-treated (**C**) cells. Volcano plots of eleven genes that support enzalutamide resistance. **D**. Most supporters of enzalutamide resistance are amplified in metastatic castration-resistant prostate cancer. Oncoprint of 429 patients from Abdia et al[16] in cBioPortal[14, 15] evaluating frequency of amplifications, deletions, and mutations. **E/F**. AR does not strongly regulate ACAT1, MAP3K11, or PSMD12. Knockdown of *AR* for three or six days in C4-2B (**E**) and 22Rv1 (**F**) cells.

Our first step was to evaluate whether our 11 genes were expressed in castration-resistant prostate cancer. First, we used cBioPortal[14, 15] to interrogate a set of publically available data from metastatic castration-resistant prostate cancer patient samples[16] with outcome and gene expression data. Unsurprisingly, *AR* was the most frequently altered of these genes, with 59% of samples bearing amplifications and mutations (Figure 1D). For the other genes, alterations were rarer, occurring in 2.8-6% of cases, depending on the gene. With the exception of *ACAT1*, which has more deletions, most of our gene list alterations were amplifications.

We also examined whether our putative resistance supporter genes increased after abiraterone and enzalutamide treatment. For this analysis, we included patients who were exposed or naïve to abiraterone and enzalutamide, and excluded patients on treatment or whose exposure status was unknown from Abida *et al*[16]. Through this analysis we determined that *AR* expression was dramatically increased in patients who had been exposed to abiraterone and enzalutamide (Supplemental Figure 1), as reported [16]. However, there was no difference in gene expression of our putative resistance drivers in patients who had received abiraterone and enzalutamide versus patients who had not (Supplemental Figure 1). With the exception of *AR*, all of our genes exhibited gains and losses in both patient groups (Supplemental Figure 2, 3).

To validate our putative supporters of enzalutamide resistance *in vitro*, we turned to siRNA knockdown studies. To complement validation studies in C4-2B cells, we also used 22Rv1 cells, which express AR-V, including AR-V7 and AR-V9, and are enzalutamide resistant[20, 21]. We used two siRNA constructs per gene and evaluated how knockdown of each gene impacts enzalutamide sensitivity and full length AR (AR-FL) and AR-V7 levels. Through this analysis, we validated three candidate genes (*ACAT1, MAP3K11*, and *PSMD12*) in two cell lines as supporters of enzalutamide resistance *in vitro*.

In our validation studies, we first assessed whether AR regulated expression of our genes of interest (Supplemental Figure 4). Our two AR constructs both efficiently knocked down expression of AR, with construct #2 being more efficient (Figure 1 E, F and Supplemental Figure 4A). Importantly, both of our knockdown constructs targeted AR-V7. Although AR-V7 expression has been reported in C4-2B cells[22], under our conditions, the expression was minimal. We next evaluated the consequences of *AR* knockdown on our genes of interest. Knockdown of *AR* reduced gene expression of *MAP3K11*, but it did not reduce the expression of *ACAT1* or *PSMD12* (Supplemental Figure 4B, C, D). We also evaluated whether knockdown of *AR* altered protein expression of ACAT1, MAP3K11, and PSMD12, and observed no significant changes. Interestingly, on a gene expression level, knockdown of many of our genes resulted in dramatic changes in expression of other members of our gene list (Supplemental Figure 4).

Our first gene of interest, *ACAT1*, encodes Acetyl-CoA Acetyltransferase 1 (ACAT1). While gene expression for *ACAT1* does not increase between benign and primary prostate cancer, *ACAT1* expression decreases in metastatic castration-resistant prostate cancer versus primary prostate cancer (Figure 2A-C). While *ACAT1* message appears to decrease between primary and metastatic castration-resistant prostate cancer, on a protein level, ACAT1 expression reportedly increases[23, 24]. For *ACAT1*, both siACAT1 constructs were efficient, and knockdown of *ACAT1* did not dramatically alter AR or AR-V7 protein expression (Figure 2D, E). Over a six-day treatment time course, knockdown of *ACAT1* resulted in a dramatic reduction in cell survival in C4-2B cells (55-76% siACAT1 enzalutamide treated versus non-targeting enzalutamide treated cells) (Figure 2F). In 22Rv1 cells, this effect was more modest; achieving a 14% survival reduction versus non-targeting enzalutamide treated cells (Figure 2G). To evaluate how ACAT1 could be supporting enzalutamide resistance, we again turned to the Abida *et al* data set[16]. Using cBioPortal[14, 15], we selected metastatic castration-resistant prostate cancer patients who were naïve to abiraterone and enzalutamide treatment and compared their AR activity and neuroendocrine (NE/NEPCa) activity scores. In clinical samples, *ACAT1* expression correlates with the AR activity score (Figure 2H) but not the NE/NEPCa score (Figure 2I).

**Figure 2:**
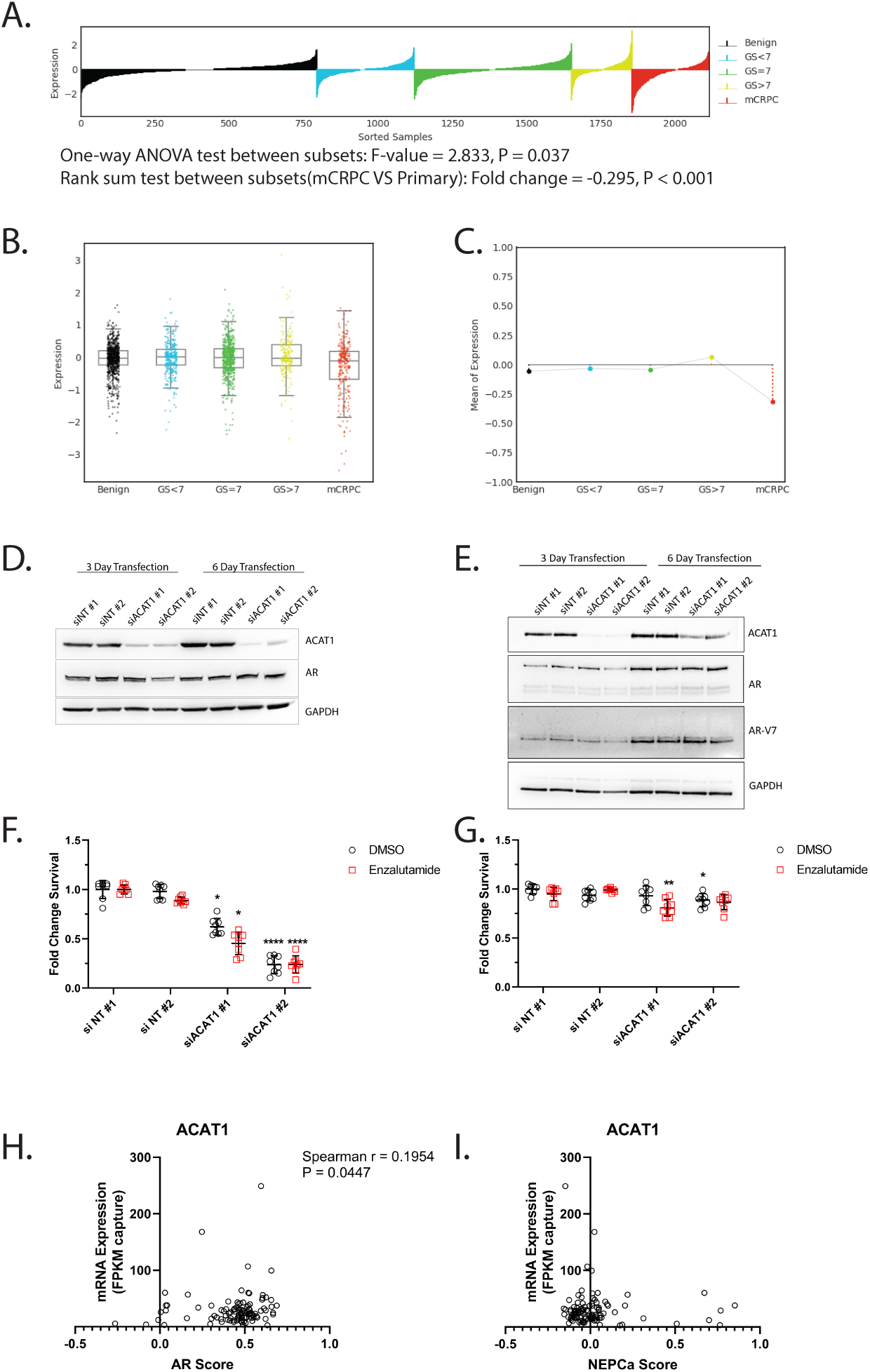
ACAT1 supports enzalutamide resistance in vitro. **A/B/C**. *ACAT1* expression decreases during progression to metastatic castration-resistant prostate cancer. Lollipop (**A**), box plot (**B**) and lineplot of mean trend (**C**) of *ACAT1* expression in benign prostate, prostate cancer, and metastatic castration-resistant prostate cancer patient samples. GS: Gleason score; mCRPC: metastatic castration-resistant prostate cancer. **D/E**. Knockdown of *ACAT1* does not strongly impact AR expression. C4-2B (**D**) and 22Rv1 (**E**) cells were transfected with one of two non-targeting siRNA (siNT) or ACAT1-targeting (siACAT1 #1 or #2) and ACAT1, AR, AR-V7 (in 22Rv1 cells) was evaluated with GAPDH used as a loading control. **F/G**. Knockdown of *ACAT1* in castration-resistant prostate cancer cells increases cell death in response to enzalutamide. C4-2B (**F**) and 22Rv1 (**G**) were transfected with one of two non-targeting siRNA (siNT) or ACAT1-targeting (siACAT1) siRNAs and challenged with 10μM DMSO (vehicle) or enzalutamide for six days. Comparisons between DMSO and DMSO, enzalutamide to enzalutamide using Kruskal-Wallis test with Dunn’s multiple comparisons test. * P < 0.05; ** P < 0.01; *** P < 0.001; ** P < 0.0001. **H**. *ACAT1* expression positively correlates with AR activity metastatic castration-resistant prostate cancer patient tissues. Correlation between *ACAT1* mRNA expression and the AR activity score evaluated by Spearman correlation in 106 abiraterone and enzalutamide naïve metastatic castration-resistant prostate cancer patients. **I**. *ACAT1* expression does not correlate with NEPCa activity metastatic castration-resistant prostate cancer patient tissues. Correlation between ACAT1 mRNA expression and the NEPCa activity score evaluated by Spearman correlation.

Our second gene of interest was *MAP3K11*, which encodes MAP3K11, also known as Mixed Lineage Kinase 3 (MLK3). *MAP3K11* expression increases dramatically during prostate cancer progression (Figure 3A-C). Both of the MAP3K11 targeting constructs knocked down *MAP3K11* expression, with construct #1 being more efficient (Figure 3D, E). The more efficient *MAP3K11* knockdown resulted in more cell death in response to enzalutamide (Figure 3F, G). In C4-2B cells, knockdown of *MAP3K11* with construct #1 and treatment with enzalutamide resulted in a 73% decrease in cell survival versus non-targeting enzalutamide treated cells, while in 22Rv1 cells this reduction was more modest with a reduction of 28%. Consistent with previous reports, when we knocked down *MAP3K11*, we observed no decrease in AR[25] or AR-V7 expression, but we did see a loss of AR-Ser650 phosphorylation in 22Rv1 cells; this decrease was more pronounced with greater MAP3K11 ablation (Figure 3E). Phosphorylation of AR-Ser650 promotes maximal AR transactivation activity[26], suggesting MAP3K11 could be supporting enzalutamide resistance through activation of AR. Interestingly, in C4-2B cells, we did not observe AR-Ser650 phosphorylation under normal cell culture conditions, consistent with previous reports where AR-Ser650 phosphorylation is induced by phorbol 12-myristate 13-acetate (PMA)[25]. In the abiraterone and enzalutamide naïve patients, *MAP3K11* expression was not statistically correlated with the AR activity score (Figure 3H), but it was negatively correlated with the NE/NEPCA score (Figure 3I).

**Figure 3:**
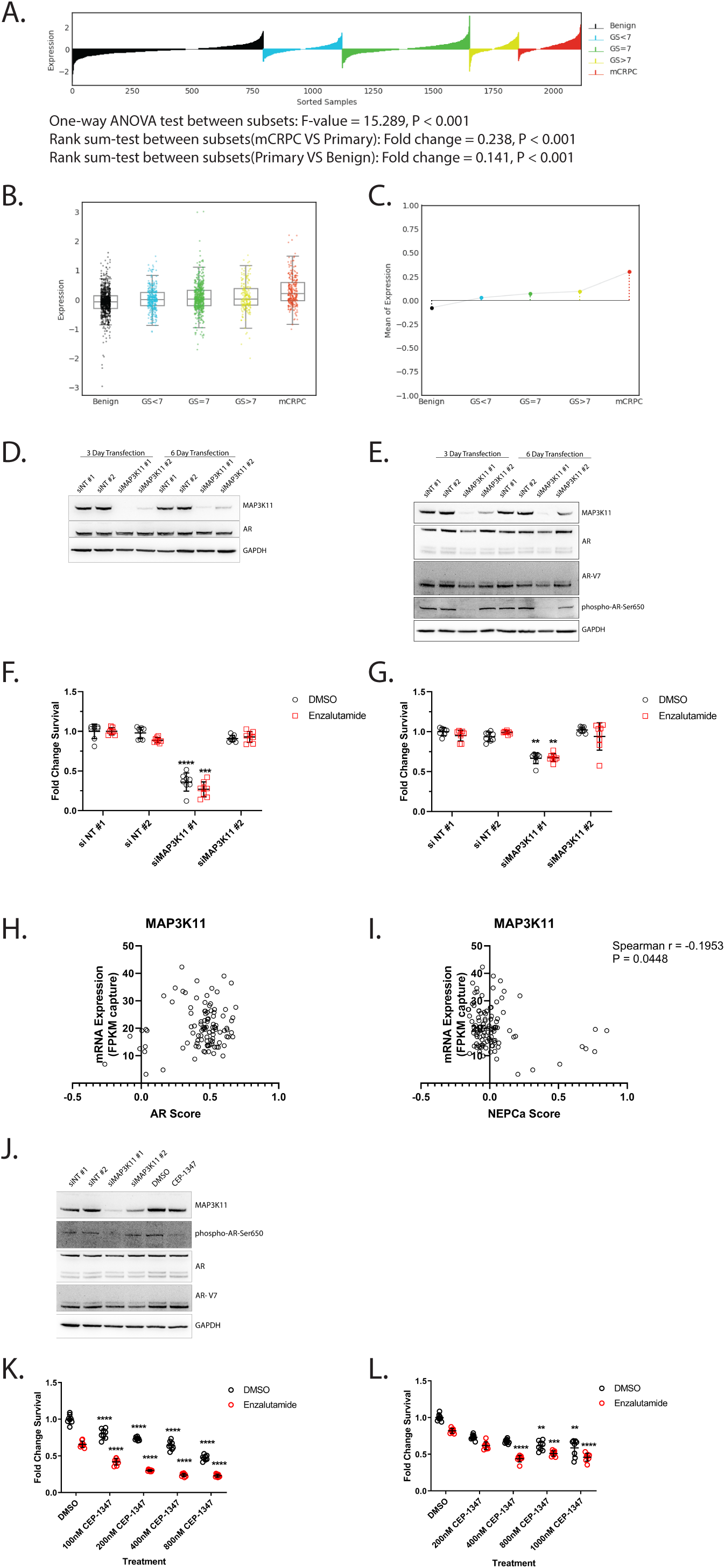
MAP3K11 supports enzalutamide resistance in vitro. **A/B/C**. MAP3K11 expression increases during progression to metastatic castration-resistant prostate cancer. Lollipop (**A**), box plot (**B**) and lineplot of mean trend (**C**) of *MAP3K11* expression in benign prostate, prostate cancer, and metastatic castration-resistant prostate cancer patient samples. GS: Gleason score; mCRPC: metastatic castration-resistant prostate cancer. **D/ E**. Knockdown of *MAP3K11* does not strongly impact AR expression but it does reduce AR-Serine 650 (AR-Ser650) phosphorylation. C4-2B (**D**) and 22Rv1 (**E**) cells were transfected with one of two non-targeting siRNA (siNT) or MAP3K11-targeting (siMAP3K11 #1 or #2) and ACAT1, AR, AR-V7 (in 22Rv1 cells), and phosphorylated AR-Ser650 was evaluated with GAPDH used as a loading control. **F/G**. Knockdown of MAP3K11 in castration-resistant prostate cancer cells increases cell death in response to enzalutamide. C4-2B (**F**) and 22Rv1 (**G**) were transfected with one of two non-targeting siRNA (siNT) or MAP3K11-targeting (siMAP3K11) siRNAs and challenged with 10μM DMSO (vehicle) or enzalutamide for six days. Comparisons between DMSO and DMSO, enzalutamide to enzalutamide using Kruskal-Wallis test with Dunn’s multiple comparisons test. * P < 0.05; ** P < 0.01; *** P < 0.001; **** P < 0.0001. **H**. MAP3K11 expression does not correlate with AR activity metastatic castration-resistant prostate cancer patient tissues. Correlation between *MAP3K11* mRNA expression and the AR activity score evaluated by Spearman correlation in 106 abiraterone and enzalutamide naïve metastatic castration-resistant prostate cancer patients. **I**. *MAP3K11* expression inversely correlates with NEPCa activity metastatic castration-resistant prostate cancer patient tissues. Correlation between *MAP3K11* mRNA expression and the NEPCa activity score evaluated by Spearman correlation. J. Knockdown or inhibition of MAP3K11 reduces AR-Ser650 phosphorylation. Comparison of AR-Ser650 phosphorylation in 22Rv1 cells transfected with non-targeting siRNA (siNT) or MAP3K11 targeting siRNA (siMAP3K11) to vector or MAP3K11 inhibitor (CEP-1347) treated cells. Importantly, AR and AR-V levels do not change. **K./L**. Inhibition of MAP3K11 with CEP-1347 potentiates enzalutamide treatment. Treatment of C4-2B (**K**) and 22Rv1 (**L**) with increasing concentrations of CEP-1347 with 10μM DMSO (vehicle) or enzalutamide. Analysis by one-way ANOVA with Sidak’s multiple comparisons test comparing 0nM CEP-1347 (DMSO) to varying CEP-1347 concentrations (black to black) and 0nM CEP-1347 (DMSO) plus enzalutamide to varying CEP-1347 concentrations plus enzalutamide (red to red). * P < 0.05; ** P < 0.01; *** P < 0.001; **** P < 0.0001

As several inhibitors have been developed for the MLK family, we next evaluated whether one of these could be used in combination with enzalutamide. CEP-1347 was initially developed to prevent HIV-1 Associated Neurocognitive Disorders (HAND) and Parkinson’s[27-29]. Treatment of 22Rv1 cells with CEP-1347 did not reduce AR or AR-V7 expression, but it did reduce AR-Ser650 phosphorylation to similar levels as *MAP3K11* knockdown (Figure 3J). Indeed, combination therapy with increasing concentrations of CEP-1347 induced increased cell death in response to 10µM enzalutamide versus vehicle treated cells in C4-2B and 22Rv1 cells in a dose-dependent manner (Figure 3K, L).

Our final gene is *PSMD12*, which encodes Proteasome 26S Subunit, Non-ATPase 12, a component of the 26S Proteasome. *PSMD12* expression does not change between benign, primary, and metastatic prostate cancer (Figure 4A-C). In our validation studies, both constructs were highly efficient at knocking down *PSMD12* gene expression in both cell lines (Figure 4D, E). In cell survival assays, we observed that C4-2B cells were exquisitely sensitive to *PSMD12* knockdown, with knockdown driving an 85-91% decrease in cell survival in C4-2B cells and a 28% reduction in survival in 22Rv1 cells (Figure 4F, G). This increased sensitivity was likely due to a loss of AR and AR-V7 expression, as when PSMD12 expression decreased, AR and AR-V7 expression decreased (Figure 4C, D). In the clinical data *PSMD12* expression was not correlated with AR activity or NE/NEPCa activity scores (Figure 4E, F).

**Figure 4:**
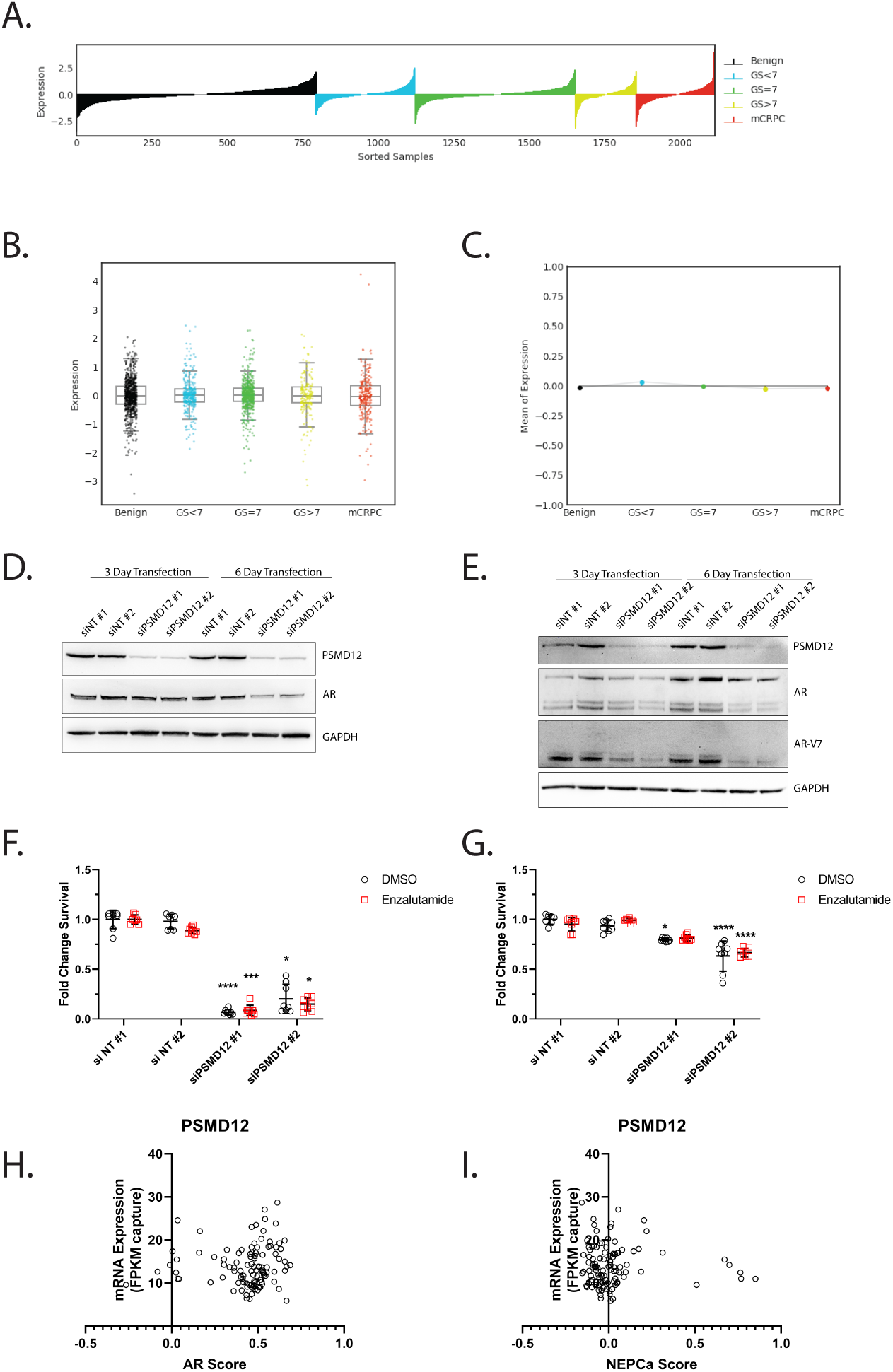
PSMD12 supports enzalutamide resistance in vitro. **A/B/C**. *PSMD12* expression is similar in benign, primary, and metastatic castration-resistant prostate cancer. Lollipop (**A**), box plot (**B**) and lineplot of mean trend (**C**) of *PSMD12* expression in benign prostate, prostate cancer, and metastatic castration-resistant prostate cancer patient samples. GS: Gleason score; mCRPC: metastatic castration-resistant prostate cancer. **D/E**. Knockdown of *PSMD12* decreases AR and AR-V expression. C4-2B (**D**) and 22Rv1 (**E**) cells were transfected with one of two non-targeting siRNA (siNT) or PSMD12-targeting (siPSMD12 #1 or #2) and PSMD12, AR, AR-V7 (in 22Rv1 cells) was evaluated with GAPDH used as a loading control. **F/G**. Knockdown of *PSMD12* in castration-resistant prostate cancer cells increases cell death in response to enzalutamide. C4-2B (**F**) and 22Rv1 (**G**) were transfected with one of two non-targeting siRNA (siNT) or PSMD12-targeting (siPSMD12) siRNAs and challenged with 10μM DMSO (vehicle) or enzalutamide for six days. Comparisons between DMSO and DMSO, enzalutamide to enzalutamide using Kruskal-Wallis test with Dunn’s multiple comparisons test. * P < 0.05; ** P < 0.01; *** P < 0.001; **** P < 0.0001. **H/I**. *PSMD12* expression does not correlate with AR activity or NEPCa activity in 106 abiraterone and enzalutamide naïve metastatic castration-resistant prostate cancer patient tissues. Correlation between *PSMD12* mRNA expression and the AR activity score (**H**) or NEPCa activity (**I**) evaluated by Spearman correlation.

## Discussion

Our shRNA screen in C4-2B cells identified 11 putative supporters of enzalutamide resistance (*TFAP2C, CAD, SPDEF, EIF6, GABRG2, CDC37, PSMD12, COL5A2, AR, MAP3K11*, and *ACAT1*). Our siRNA constructs efficiently targeted all of our genes of interest, and revealed that, at least on a gene expression level, there is considerable cross-talk between these genes. For example, knockdown of *MAP3K11*, results in decreased expression of genes that are regulated by AR, likely be reducing its transcriptional efficiency. Through our validation studies, we verified that transient knockdown of *ACAT1, MAP3K11*, or *PSMD12* could sensitize castration-resistant prostate cancer cells to enzalutamide.

In general, C4-2B cells were much more sensitive to knockdown of these three genes in combination with enzalutamide treatment. There are two possible explanations for this observation. The first is that we identified these genes as supporting enzalutamide resistance in C4-2B cells through our shRNA screen, and thus these cells are more dependent on these pathways. Alternatively, it is possible that the high level of AR-V in 22Rv1 cells, which confers enzalutamide resistance by removing dependency on ligand binding, simply makes 22Rv1 cells much more resistant. As knockdown of *PSMD12* reduces AR and AR-V7 expression, and 22Rv1 cells do not become dramatically more sensitive to enzalutamide, this would suggest some of these genes might only be required for enzalutamide resistance in C4-2B cells, and by extension, only a subset of prostate cancers.

One of our enzalutamide-resistance supporting genes, *ACAT1*, encodes the protein ACAT1 that has several roles in the mitochondria. First, it serves as the final enzyme in isoleucine metabolism, converting 2-methylacetoacetyl-CoA into propionyl-CoA and acetyl-CoA[30]. ACAT1 also functions in ketone body metabolism in which its substrate is acetoacetyl-CoA[30]. Produced acetyl-CoA is shuttled into the Krebs cycle to be oxidized for energy production. More recently, ACAT1 has been implicated in regulating the pyruvate dehydrogenase complex, comprised of pyruvate dehydrogenase and PDH phosphatase, through acetylation[31]. Importantly, knockdown of ACAT1 activity is sufficient to push cancer cells from their preferred aerobic glycolysis, favored by cancer and proliferating cells, and into oxidative phosphorylation, the energy production mechanism favored by differentiated cells. Beyond its role in metabolism, ACAT1 dysregulation could alter acetyl-CoA levels. Notably, AR can be acetylated at AR-K618 [32] in the DNA binding domain and AR-K632, AR-K633[33] and AR-K630[34] in the hinge region, which supports increased AR transcriptional activity and increased tumor growth in xenograft models[32, 34]. Which of these mechanisms supports enzalutamide resistance remains to be determined.

In prostate, ACAT1 is associated with more aggressive prostate cancer and castration-resistant disease[23, 24]. These observations, combined with our experimental data, suggest that ACAT1 is expressed in castration-resistant prostate cancer and may support enzalutamide resistance, making it an attractive therapeutic target. ACAT1 activity can be inhibited by arecoline hydrobromide[31] or the FDA-approved drug sulfasalazine[35]. Unfortunately, neither arecoline hydrobromide nor sulfasalazine are specific to ACAT1.

Another gene of interest *MAP3K11*, encoding MAP3K11, perhaps holds the most therapeutic potential. MAP3K11 is a serine/threonine kinase[36], which via phosphorylation of MAP2Ks (MKKs), is a regulator of the mitogen-activated protein kinases (MAPKs), including JNK, ERK, and p38 (reviewed[37]). MAP3K11 can act both as a kinase and a scaffold for other kinases[37]. Earlier studies identified MAP3K11 as a potent regulator of AR transcriptional activity in C4-2B cells through an RNAi phenotypic screen[25]. The link between MAP3K11 and AR activity appears to be driven by phosphorylation of serine 650, located in the hinge region of AR[33]. The hinge region is absent in most AR-Vs, including AR-V7, but it is present in a subset, including AR^v567es^ [33], suggesting this posttranslational modification could regulate the activities of some AR-Vs. Based on alanine-for-serine mutation studies using human AR in COS cells, phosphorylation of AR-Ser650 is required for optimal AR transactivation activity, as without AR-Ser650 phosphorylation, transcription of an AR-responsive mammary tumor promoter was reduced by 30%[26]. While it is possible that AR-Ser650 is a direct target of MAP3K11, *in vitro* kinase assays revealed AR-Ser650 is phosphorylated by JNK and p38 [38], which are downstream of MAP3K11. Which of these pathways are effectors of MAP3K11 in castration-resistant prostate cancer is currently under investigation.

Of our three genes, MAP3K11 has the most specific and tested therapeutics. Importantly, there are several inhibitors that are somewhat selective for MAP3K11 versus other mixed lineage kinase family members: CEP-1347 and URMC-099. CEP-1347 was initially developed to prevent HAND and Parkinson’s[27-29], and while safe and well-tolerated it failed to prevent Parkinson’s progression in a Phase II clinical trial[39]. CEP-1347 is a fairly selective MAP3K11 inhibitor (IC_50_ = 23 nM[28]; IC_50_ = 6 nM[40]), with limited off-target effects on other MLK family members (MLK1 IC_50_ = 38 nM and MLK2 IC_50_ = 51 nM[28]; MLK1 IC_50_ < 1 nM and MLK2 IC_50_ = 2 nM[40]). URMC-099 is a more specific MAP3K11 inhibitor (IC_50_ = 14 nM), with limited off-target effects on other MLK family members (MLK1 IC_50_ = 19 nM; MLK2 IC_50_ = 42 nM; and DLK IC_50_ = 150 nM)[40]. Our current studies are focusing on evaluating these compounds as a therapy in conjunction with enzalutamide in enzalutamide-resistant castration-resistant prostate cancer.

Our final gene, *PSMD12*, has not been previously implicated in prostate cancer, but dysregulation of the proteasome has been associated with prostate cancer. Inhibition of the proteasome with therapeutics like bortezomib has been shown to increase the efficacy of first generation anti-androgens like bicalutamide by decreasing the expression of AR and AR-V [41] or induce sensitivity of AR-independent prostate cancer cells to etoposide[42]. Unfortunately, proteasome inhibitors, to date, have proven acutely toxic, and targeting PSMD12 with currently available therapeutics is unlikely to provide a favorable risk-to-benefit ratio.

Although not all of our candidate genes validated in our follow-up studies, it should be noted that many of these hold promise and may be better evaluated in other cell lines and assays. For example, *GABRG2* encodes the Gamma-Aminobutyric Acid Type A Receptor Gamma2 Subunit protein, which is a GABA receptor. Previous studies by Jin *et al* discovered *GABRG2* as a member of a prognostic 21-gene panel (NARP21) that predicted decreased overall cancer-specific survival and metastasis-free survival of patients with prostate cancer [43]. Similarly, *CAD*, which encodes carbamoyl-phosphate synthetase II, aspartate transcarbamylase, and dihydroorotase is associated with the synthesis of pyrimidine nucleotides, necessary for cell proliferation. CAD is regulated by the MAPK cascade[44], and indeed knockdown of *MAP3K11* reduces *CAD* gene expression. CAD also interacts with AR and fosters AR translocation into the nucleus, and is posited as an early marker of prostate tumor recurrence[45]. Finally CDC37, which directs kinases to the HSP90 complex, has been implicated in supporting prostate cancer cell growth via increasing AR activity and activation of kinases [46]. The failure to validate some of these potential supporters of enzalutamide resistance, therefore, is likely to be due to the limitations of our assay rather than their lack of importance in prostate cancer biology.

These studies have several limitations. First, we have focused on genes that are already expressed in castration-resistant prostate cancer cell lines rather than those that have emerged because of treatment with enzalutamide. We have also only focused on castration-resistant prostate cancer treated with enzalutamide, as at the initiation of our studies, enzalutamide treatment was not yet approved in the metastatic hormone-sensitive setting[47]. In addition, our study has been limited to 5,043 genes that are involved in signal transduction and are drug targets, which has left out a considerable number of potential drivers, like transcription factors, that likely play an important role in enzalutamide resistance. We have also performed our validation experiments in the context of transient knockdown experiments *in vitro* and focused exclusively on AR-centric resistance mechanisms. It is likely that ACAT1, MAP3K11, and PSMD12 act beyond AR to support enzalutamide resistance. Subsequent studies will focus on defining these enzalutamide-resistance drivers more holistically, including both loss of function and gain of function experiments to delineate how these genes support enzalutamide resistance both *in vitro* and *in vivo*. In summary, our studies have identified 11 genes (*TFAP2C, CAD, SPDEF, EIF6, GABRG2, CDC37, PSMD12, COL5A2, AR, MAP3K11*, and *ACAT1*) that support enzalutamide resistance in castration-resistant C4-2B cells, and we have validated three of these genes in enzalutamide-resistance *in vitro* (*ACAT1, MAP3K11*, and *PSMD12)*.

## Acknowledgements

We would like to thank Jianghong Zhang, PhD, formerly of Vanderbilt University Medical Center, for technical assistance with the shRNA screen.

---

**Supplemental Figure 1:**
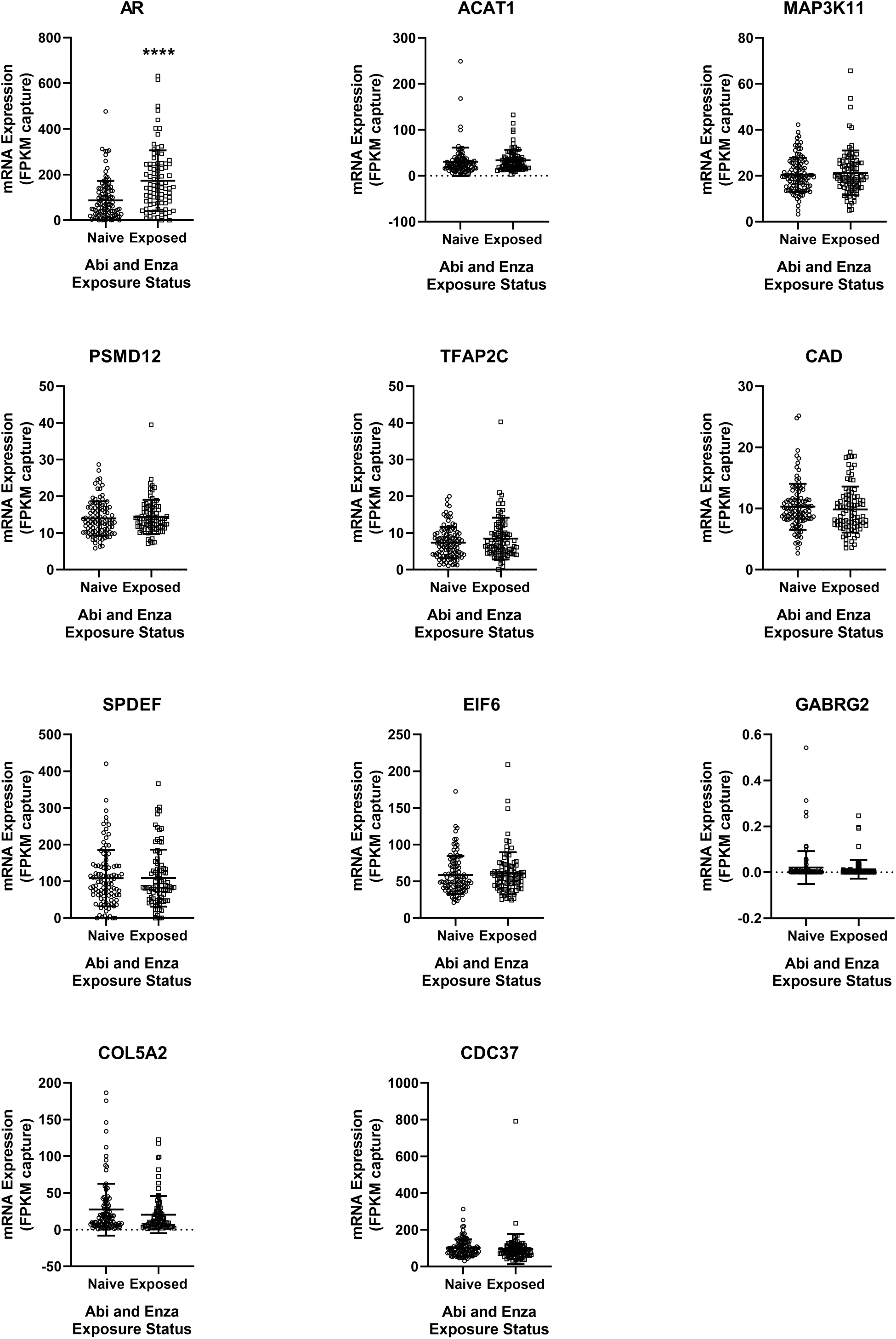
Most putative enzalutamide-resistance drivers are not increased in response to abiraterone and enzalutamide treatment in patient samples. Gene expression changes of our putative drivers in metastatic castration-resistant prostate cancer in response to abiraterone and enzalutamide. Analysis of publically available data [16] with associated outcome and gene expression data [17], using cBioPortal[14, 15] interrogated using the Mann-Whintey U test. **** P < 0.0001.

**Supplemental Figure 2:**
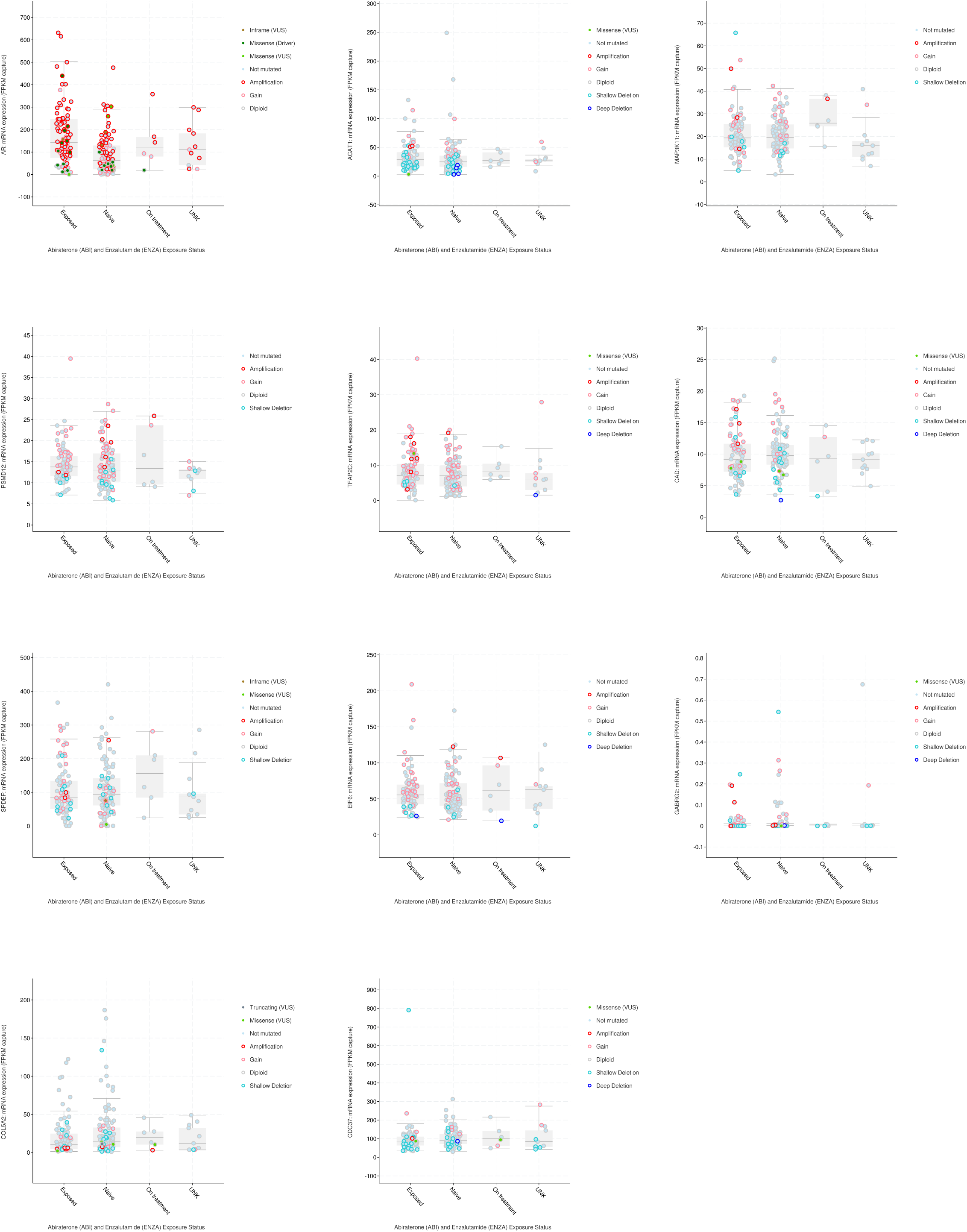
Genetic alterations in putative resistance drivers based on abiraterone and enzalutamide status. Gene expression changes overlapped with gene mutations/amplifications/deletions of our putative drivers in metastatic castration-resistant prostate cancer in response to abiraterone and enzalutamide. Analysis of publically available data [16] with associated outcome and gene expression data [17], using cBioPortal[14, 15]. VUS: variants of unknown significance

**Supplemental Figure 3:**
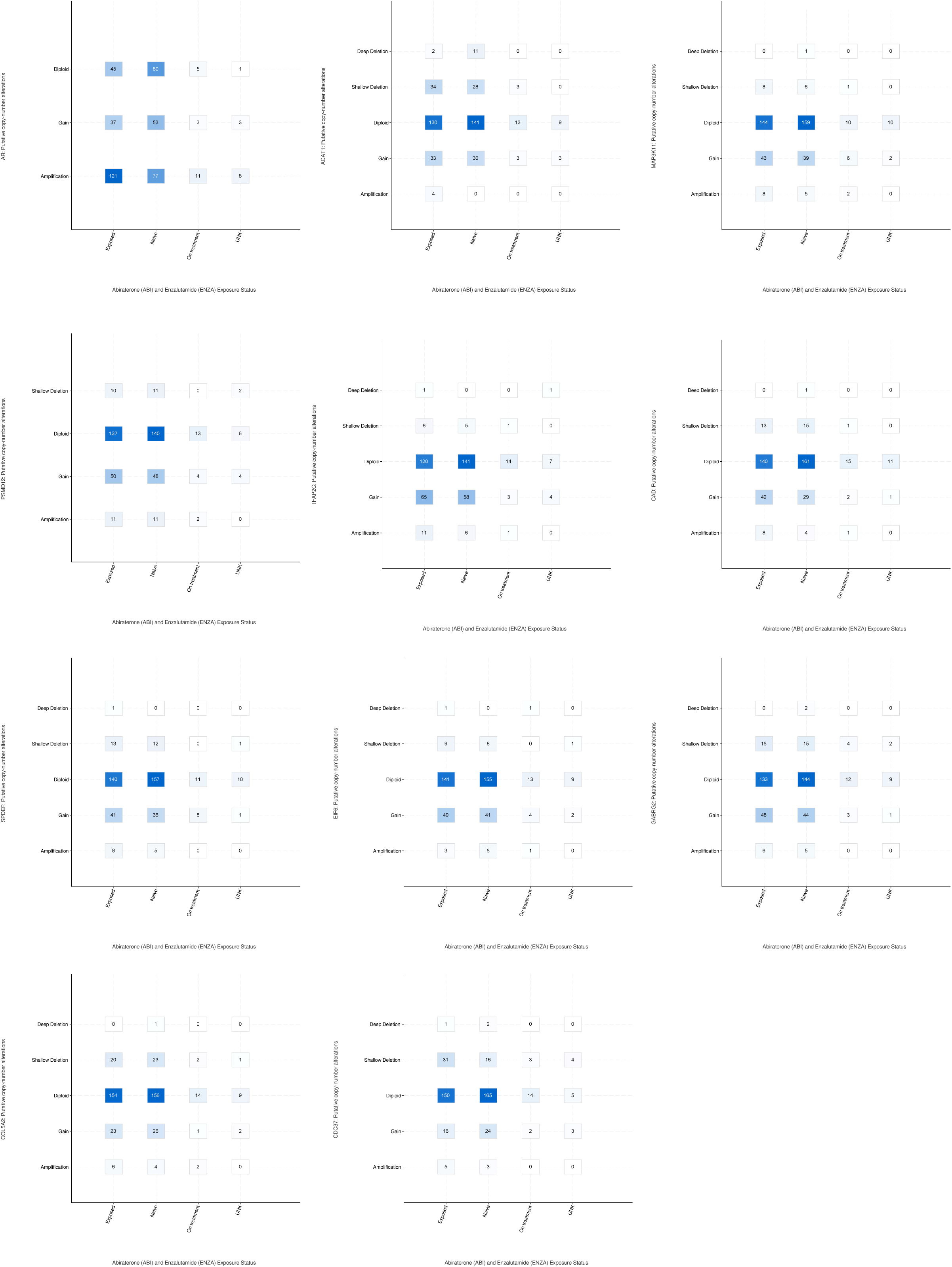
Breakdown of genetic alterations by category in putative resistance drivers based on abiraterone and enzalutamide status. Gene alterations of our putative drivers in metastatic castration-resistant prostate cancer in response to abiraterone and enzalutamide. Analysis of publically available data [16] with associated outcome and gene expression data [17] using cBioPortal[14, 15].

**Supplemental Figure 4:**
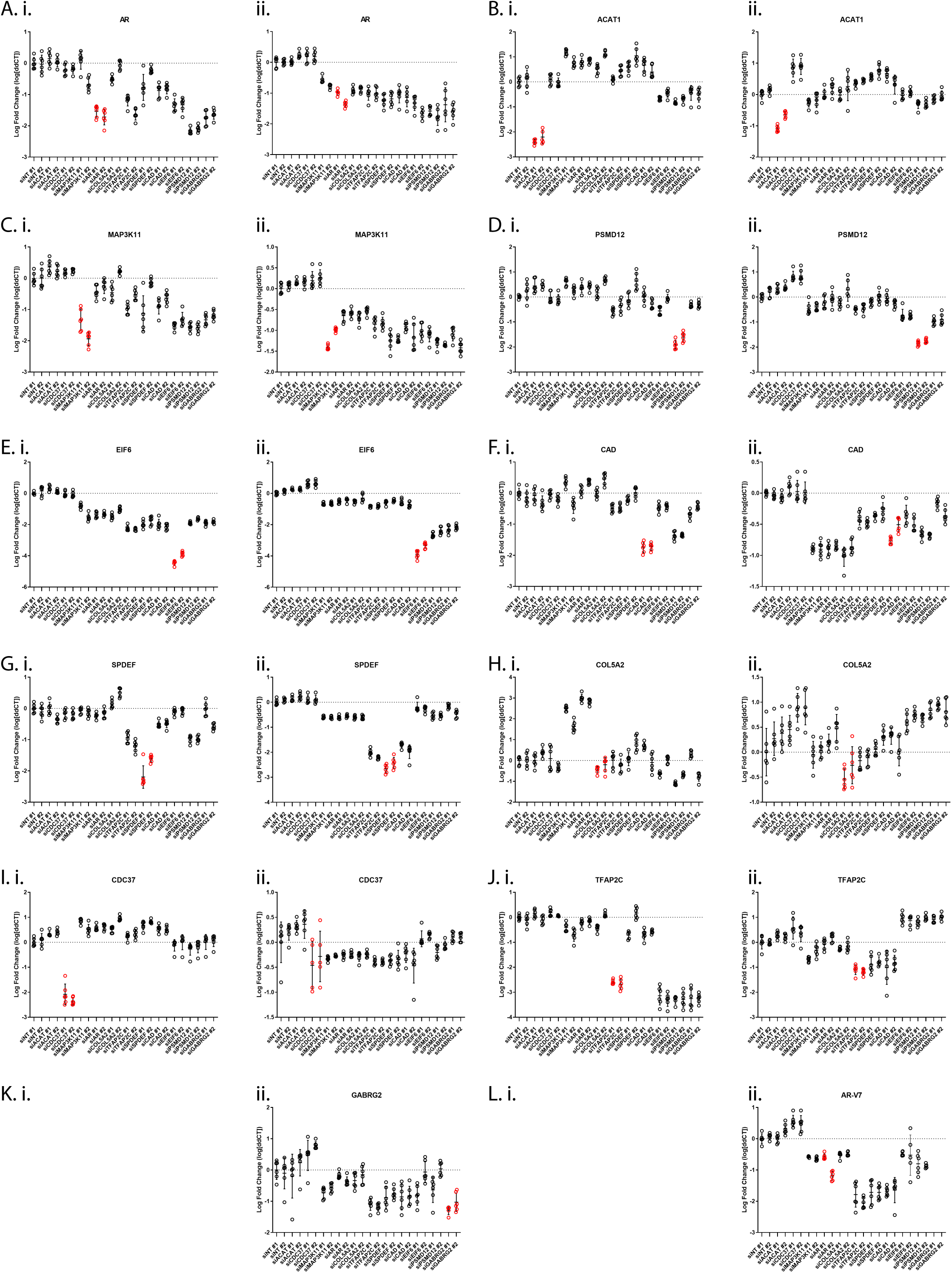
Verification of knockdown of enzalutamide-resistance drivers. Knockdown of our eleven target genes (A-L) in C4-2B (i) and 22Rv1 (ii) cells. C4-2B cells do not express *GABRG2* or *AR-V7* at sufficient levels for detection. Data presented at log fold change.

## Supplemental Methods To

## Introduction

In 2012, the US Food and Drug Administration approved enzalutamide (Xtandi) for the treatment of patients with metastatic castration-resistant prostate cancer who have previously received docetaxel. In most patients, enzalutamide halts progression of the disease for a short period of time, but patients usually relapse, which is associated with poor prognosis. Identifying the genes and pathways that the cancer cells use to overcome enzalutamide are of interest, because new interventions aimed at preventing relapse after enzalutamide therapy would have therapeutic value to late-stage prostate cancer patients.

In this study, we took advantage of RNA interference techniques and an *in vitro* model of castration-resistant prostate cancer cells to identify genes that drive resistance to enzalutamide. We transduced one-third of a commercially available bar-coded lentiviral shRNA library (DECIPHER from Cellecta) into C4-2B cells. The shRNA library contained shRNA clones targeting approximately 5045 human genes. The shRNA integrated into the chromosomal DNA of the transduced C4-2B cells and stably knocked down messenger RNA transcripts.

Our methodology called for the establishment of three populations of transduced C4-2B cells, which were maintained for six days before the cells were harvested and genomic DNA extracted. The first population (hereafter, “Initial”) is a control that was maintained in the absence of enzalutamide and DMSO, which is the solvent for enzalutamide. The second population (“EnzNeg”) was maintained in the presence of DMSO, but not enzalutamide, to control for the effects of the solvent. The third population (“EnzPos”) was a test population maintained in the presence of both DMSO and enzalutamide.

Genomic DNA from 2 biological replicates from Initial and 3 biological replicates from each of EnzNeg and EnzPos were subjected to targeted deep sequencing on an Illumina instrument (total of eight samples). The targeting protocol promoted specifically the sequencing of the bar-coded shRNA in the genome.

In this analysis, we count the number of times each shRNA bar-code was sequenced in the eight samples, and calculate the ratio of the abundance of each shRNA in EnzPos versus both control groups. Our hypothesis is that if a shRNA-targeted gene helps cancer cells to achieve resistance when selected by enalutamide, we will observe fewer bar-codes for those genes relative to both controls (which translates into more mRNA for the gene in EnzPos than both controls). A given shRNA was considered a “hit” if it showed at least two-fold abundance decrease in enzalutamide-positive relative to both enzalutamide-negative control and initial samples.

## 1 Report Linux Shell Variables

This first portion of the shRNA data analysis is coded in Linux bourne-again shell using freely available command-line tools. We declared the following directories as shell variables to condense and simplify the code for this analysis. The variables’ paths were set in knitR’s setup chunk using `Sys.setenv()`:

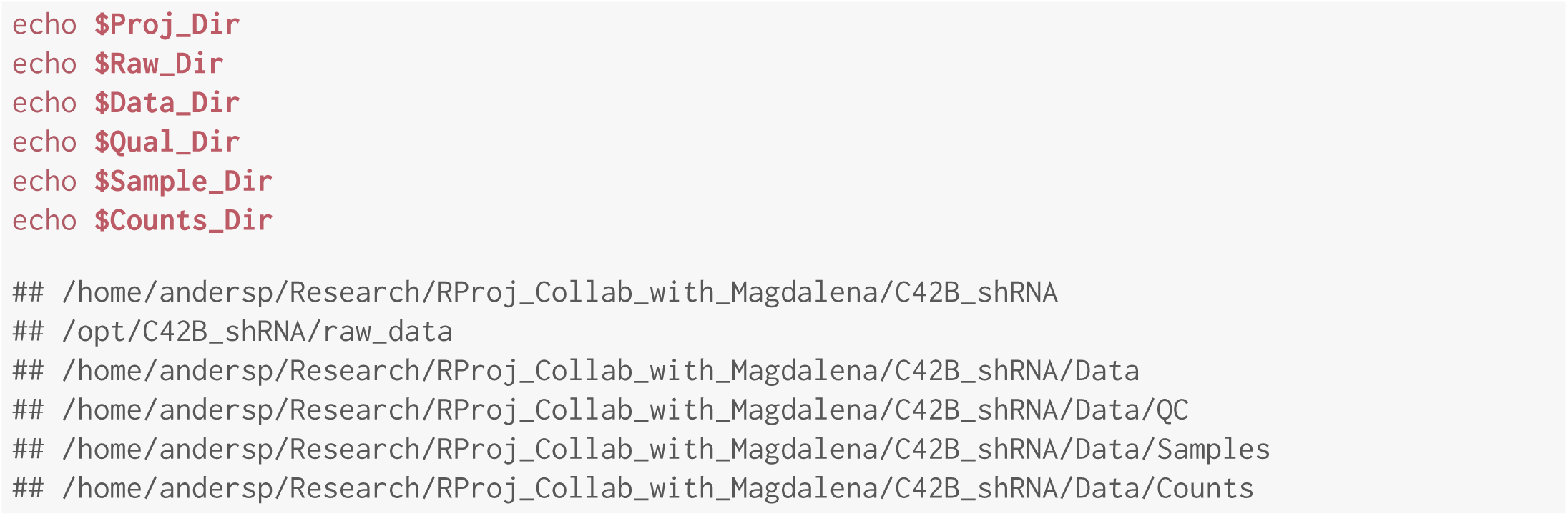

## 2 Sample Preparation

Here we extract the compressed raw data using the `unzip` command from the Ubuntu Linux *utils* package. The code tests if the uncompressed data directory already exists and if so, deletes it, in keeping with the spirit of reproducible research. The uncompressed data directory is recreated by the `unzip` command without the need to invoke `mkdir`. The -j argument directs `unzip` to junk the archive’s directory structure and dump all the files into the data (-d) directory.

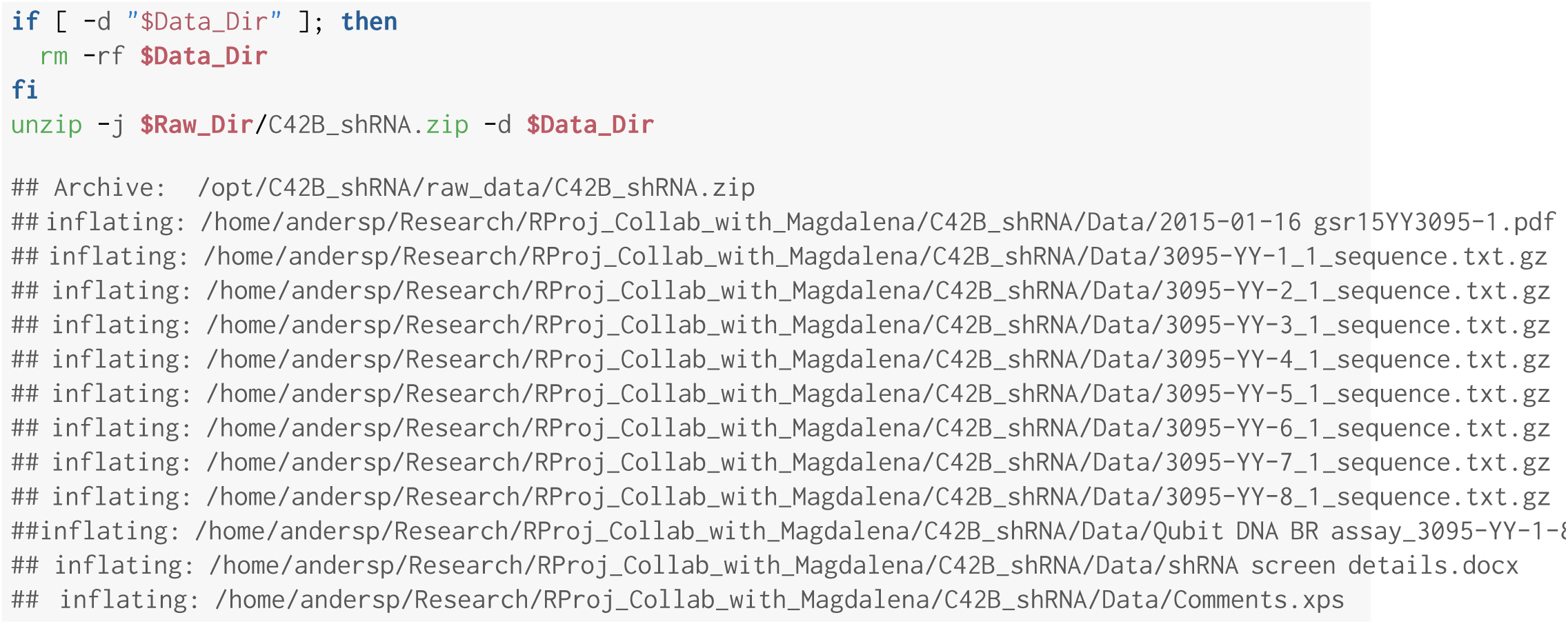

### 2.1 Quality Control

Quality control on the sequences is performed using FastQC [1]. This code runs FastQC for each of the eight sample files, and saves the results to the QC directory for us to inspect. By default, FastQC prints progress to the console, which is suppressed here with the –quiet argument. The –threads 8 argument directs FastQC to run in parallel for each of the eight samples, one CPU core each, without needing to invoke GNU parallel. Setting –threads to a number greater than eight does not make the code run faster. Linux `mkdir` is necessary because FastQC will not create the output directory (–outdir) if it does not already exist.

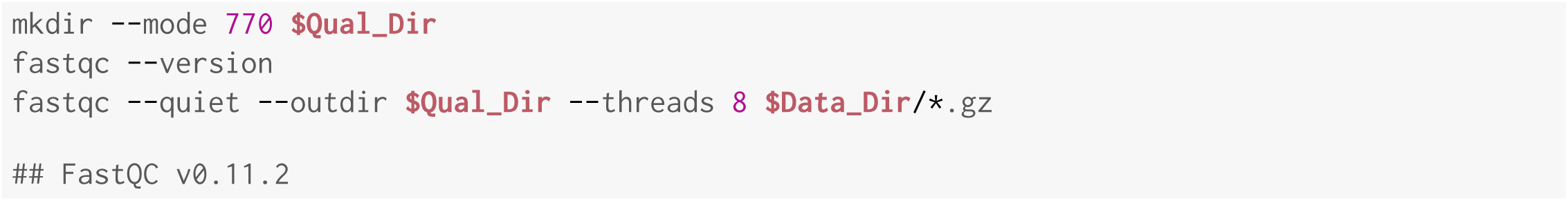

FastQC showed that the first eighteen nucleotides of the 50-nucleotide sequences were high quality. These eighteen nucleotides are the CELLECTA barcodes that are the subject of this analysis.

## 3 Analysis

To analyze the barcodes, we will employ an in-house script that retains the first eighteen nucleotides from each sequence.

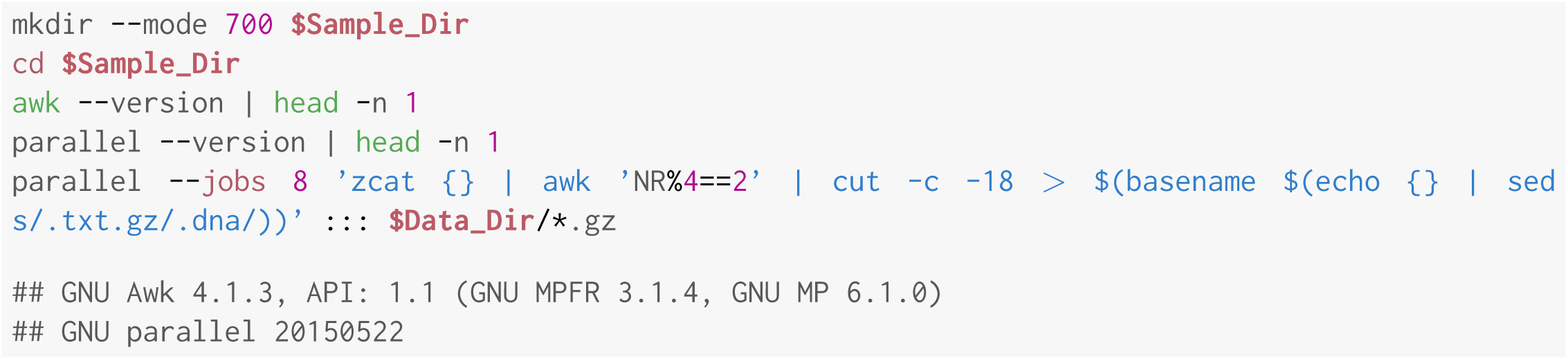

The shRNA barcodes were tallied using the table() function in R. The following code was run in an Rscript to facilitate running using GNU parallel. This code is the content of the Rscript:

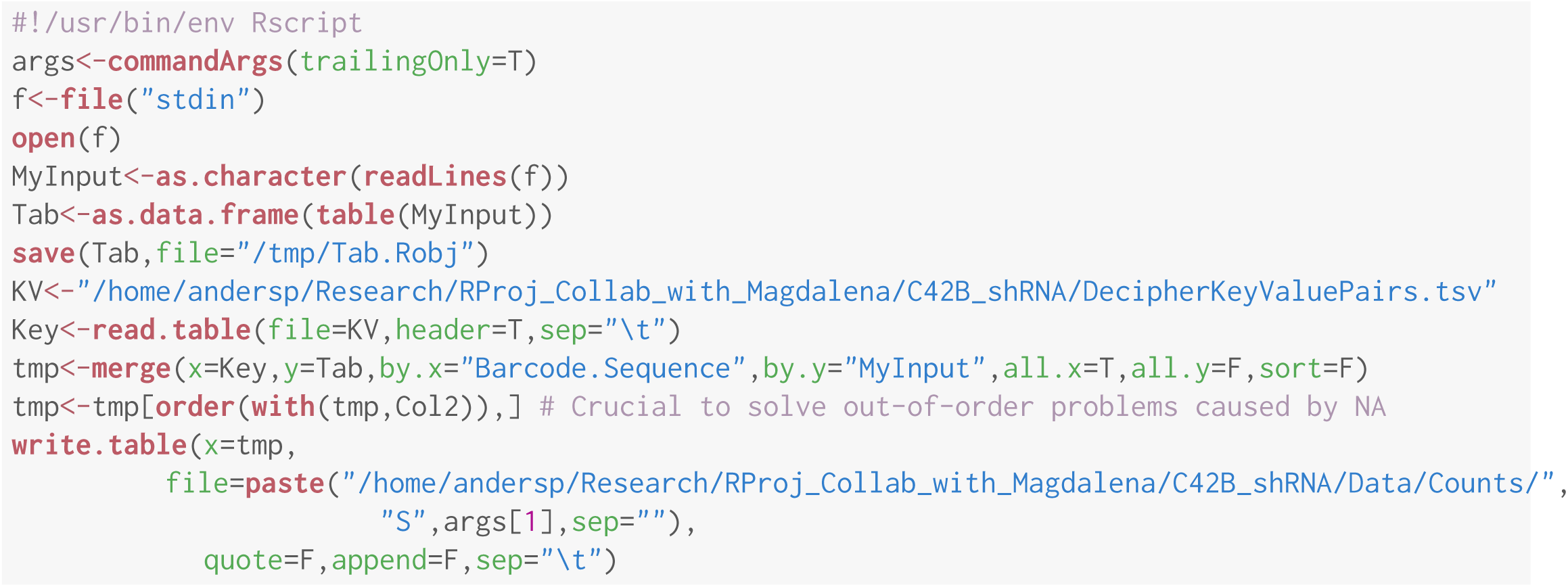

Linux inherits the Unix trait that disallows use of unexported variables in shells started from the running shell (as in bash -c). GNU parallel’s `env_parallel` argument is still in beta testing. Therefore we use the full paths in the command below, which renders it a bit ugly.

Running eight iterations of Count_shRNA.R on eight CPU cores (one core for each sample). This code takes about 45 seconds to run:

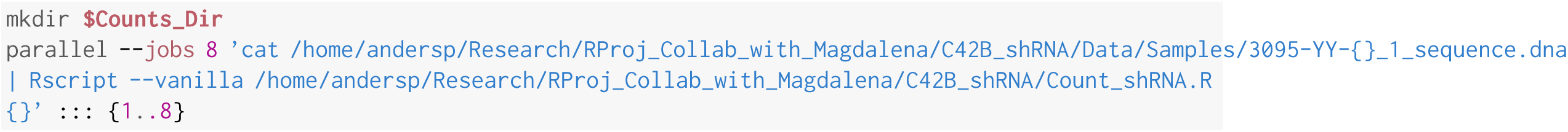

Below is an example of the results from sample 1 for the Vimentin mRNA transcript. Vimentin chosen because the gene has a short name and most of the data will fit on this page.

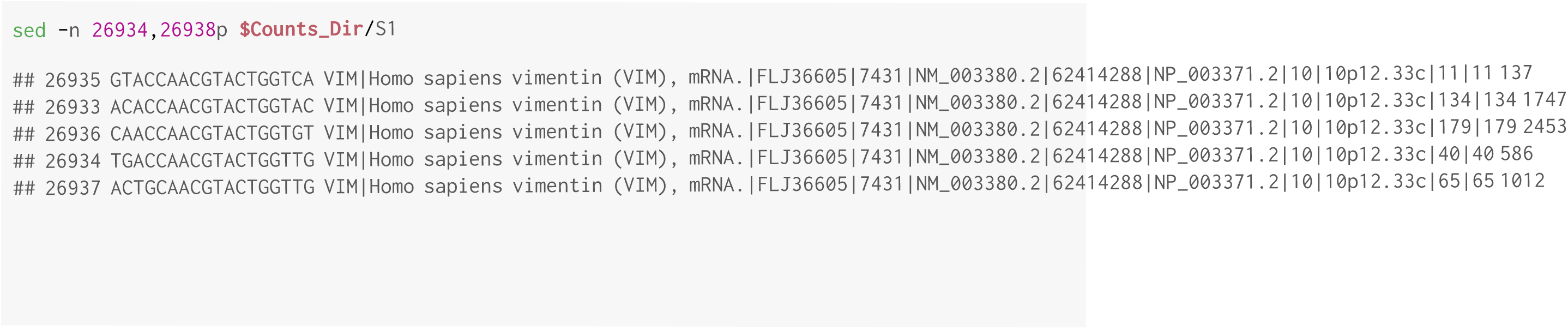

The data is arranged in four columns. The first column is an arbitrary row number that can be ignored. The second column contains the shRNA barcode against the transcript being silenced. The third column is annotation data that informs the transcript identity, NCBI database identifiers and cytoband information. The fourth column indicates the frequency at which the barcode was detected by deep sequencing the library prepared from this sample. For most transcripts being silenced, there are five barcodes from two different shRNA molecules targeting the transcript. We plan to compare barcode frequency between different samples to test the effect of enzalutamide on silencing.

The next step is to combine all the counts data into a single file:

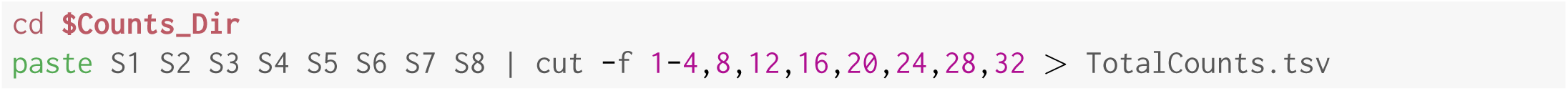

## 4 Analysis of shRNA counts in R

The remained of the workflow is coded in the R language for statistical analysis. Here we import the data into R, ignoring the header line:

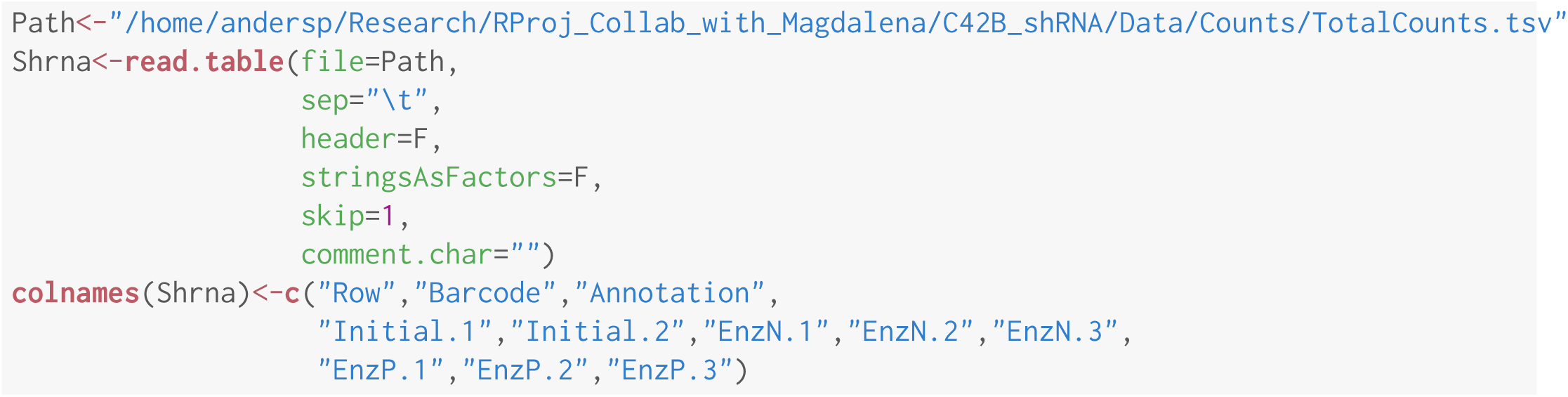

At least one sample contains NA values which are actually zeroes. A zero means that the barcode in question was never found in the raw sequencing data. Here we coerce the NAs to zeroes:

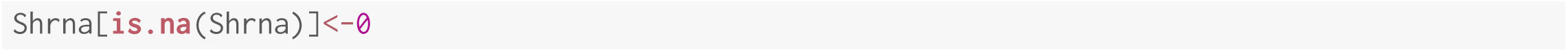

Further processing:

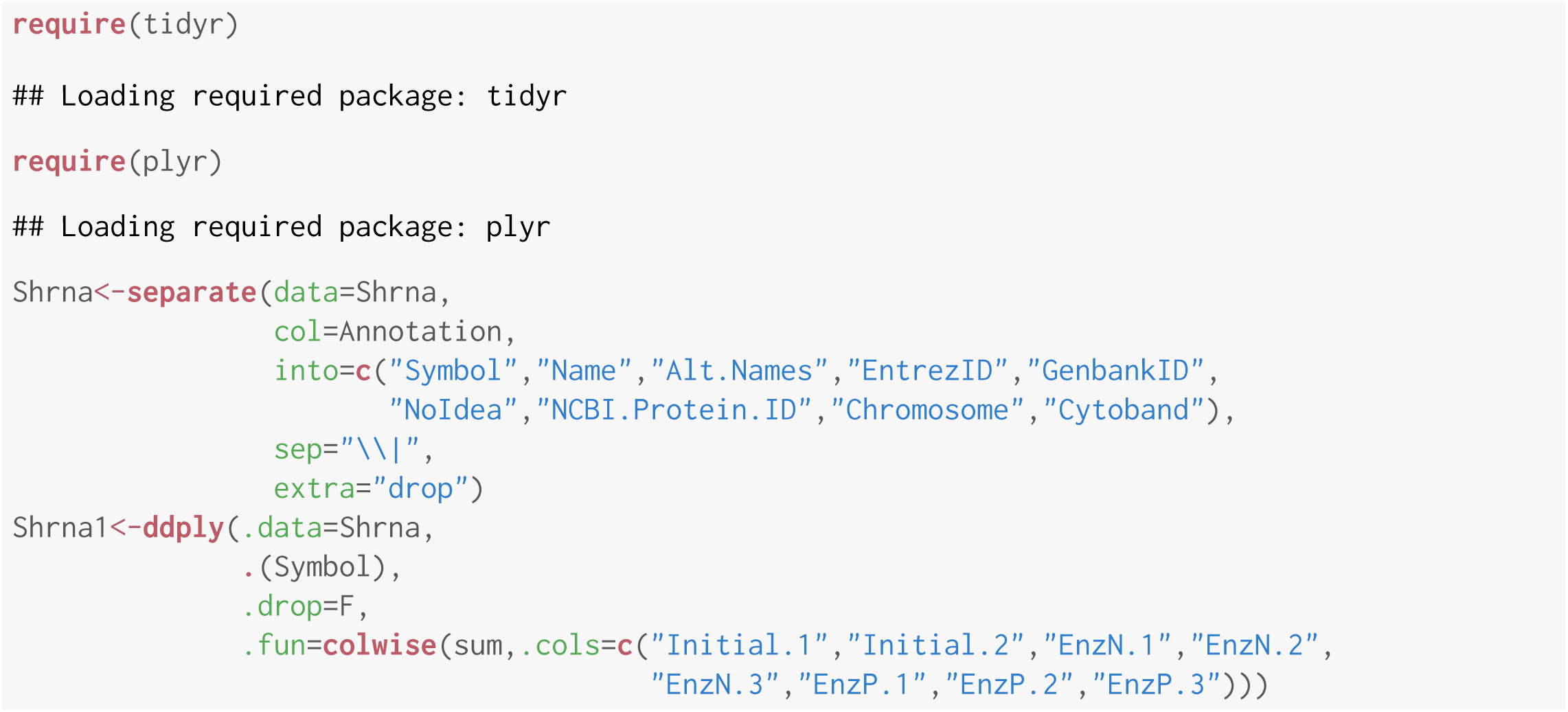

## 5 Differential Counts Analysis

EdgeR [2] and Limma [3] are R-Bioconductor packages for differential expression based on count data, which can be adapted for our purposes. Here we prepare the samples for analysis by creating a convenient object to store the count information and sample identities, called a DGEList.

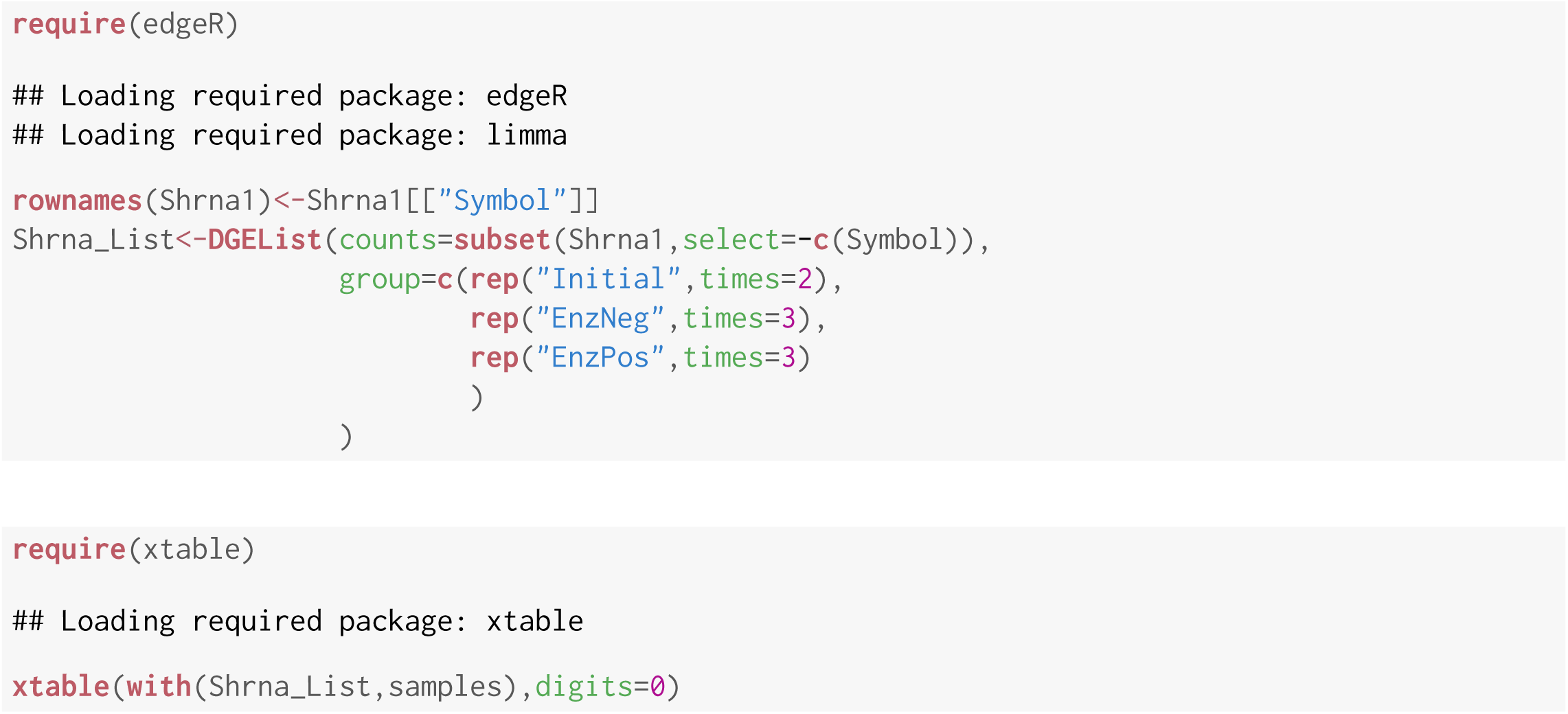

Next we apply a filter to remove genes from the analysis if they consistently have zero or very few counts. The method used in the edgeR vignette is to keep only those genes that have at least one read per million in at least 3 samples, because such low counts would make the differential counts analysis troublesome.

The following command determines the number of counts per million reads sequenced in each sample 5 (`cpm`) and tests if the number is greater than one. If it is, then a TRUE result will be produced. The `rowSums` command is a convenience that has the same functionality as `apply` by rows: it coerces the TRUE/FALSE output to one or zero, then adds all the numbers together. The maximum number that can be produced for each gene in this study is eight, which would indicate that all eight samples have at least one read per million for the gene in question. We will drop genes that are known from fewer than one read per million if that situation occurs in three or more of our eight samples.

|  | group | lib.size | norm.factors |
| --- | --- | --- | --- |
| Initial.1 | Initial | 22141545 | 1 |
| Initial.2 | Initial | 19125653 | 1 |
| EnzN.1 | EnzNeg | 20202085 | 1 |
| EnzN.2 | EnzNeg | 19523160 | 1 |
| EnzN.3 | EnzNeg | 20178264 | 1 |
| EnzP.1 | EnzPos | 21474089 | 1 |
| EnzP.2 | EnzPos | 20831135 | 1 |
| EnzP.3 | EnzPos | 22619107 | 1 |

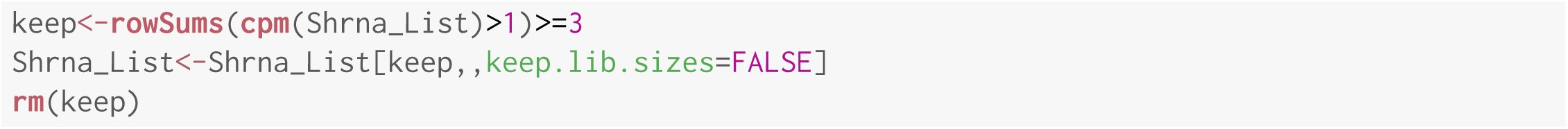

In this case, no genes were removed.

Here we apply scale normalization to the shRNA read counts, using the TMM normalization method:

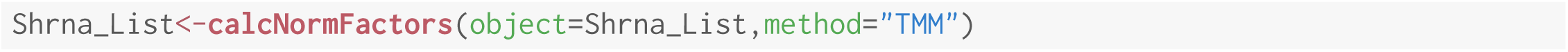

### 5.1 Differential Counts by the Limma-Trend Method

Limma-trend is a straightforward test for differential counts analysis that works well when all the samples to be analyzed have approximately the same sequencing depth. The yardstick used in the Limma User’s Guide is a ratio of the largest library size to the smallest is less than 3-fold:

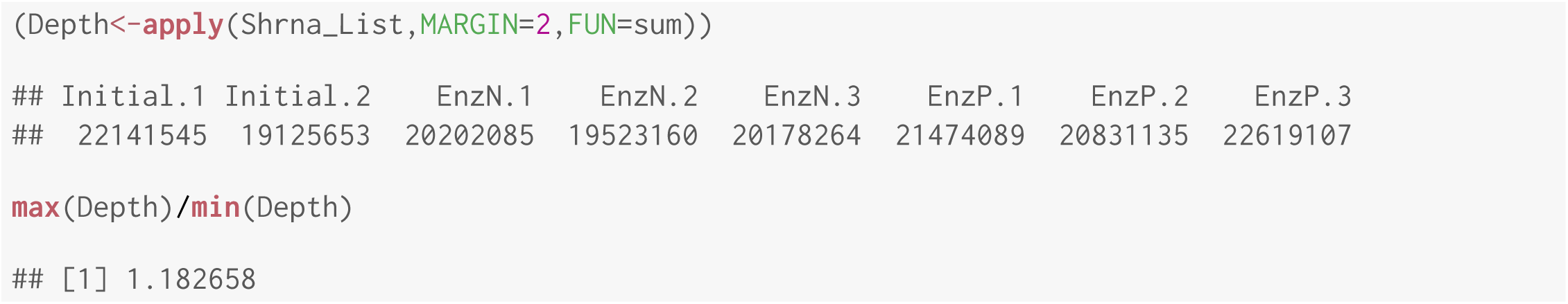

In our case, the ratio of the largest library size to the smallest is 1.182658, so the key assumption of equivalent library sizes is satisfied.

For the Limma-Trend method, counts are converted to logCPM values using edgeR’s `cpm` function. The `prior.count` is a count to be added to each observation to avoid taking the log of zero, which also reduces the variances of logarithms with low counts:

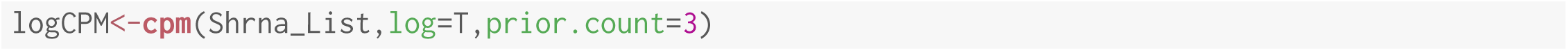

Here we make a design matrix to inform Limma of our test group (EnzPos) and both control groups (EnzNeg and Initial):

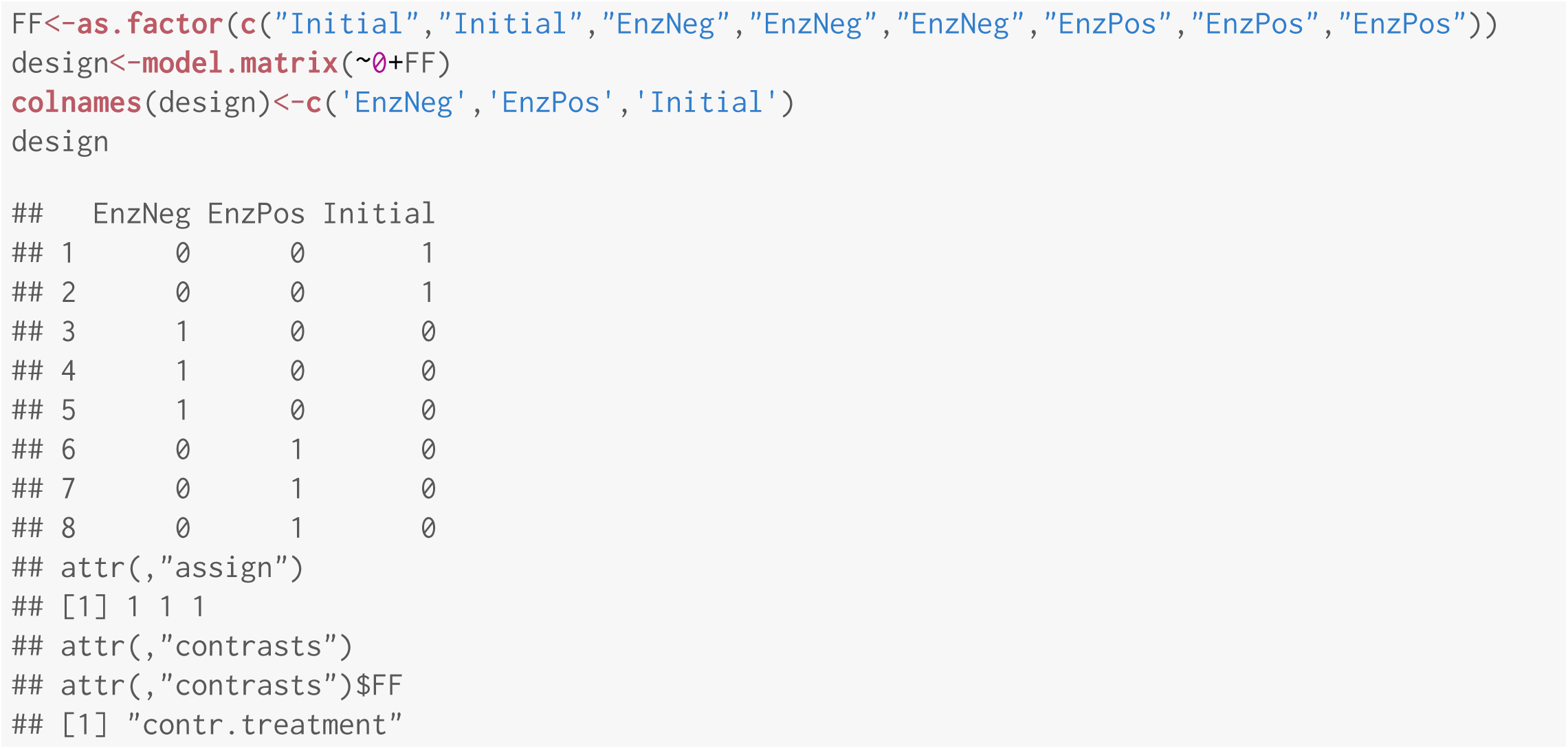

Applying the design matrix to the fitted object:

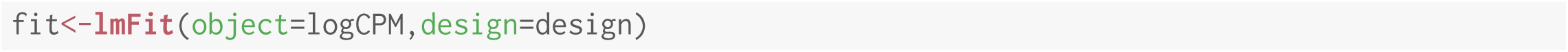

Building the contrast matrix:

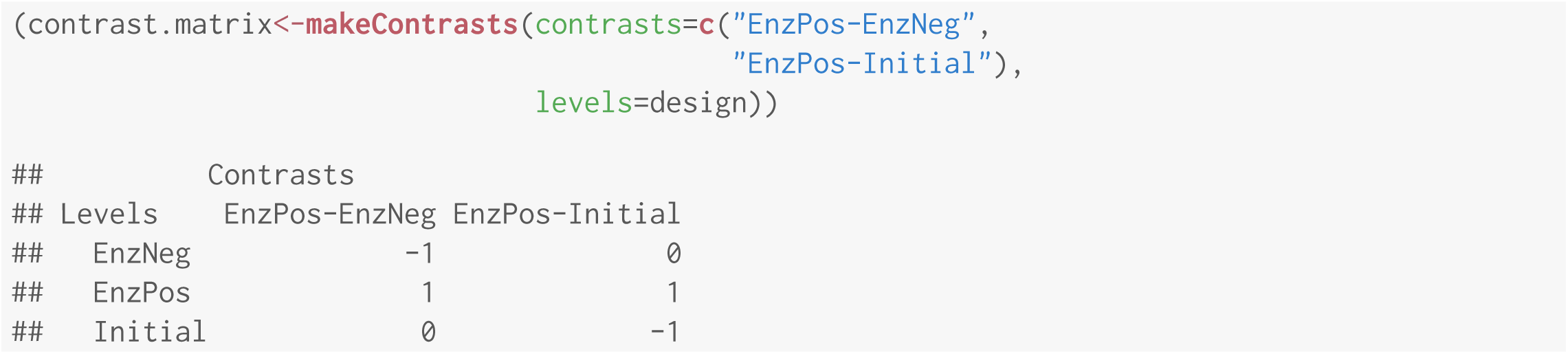

Applying the contrast matrix:

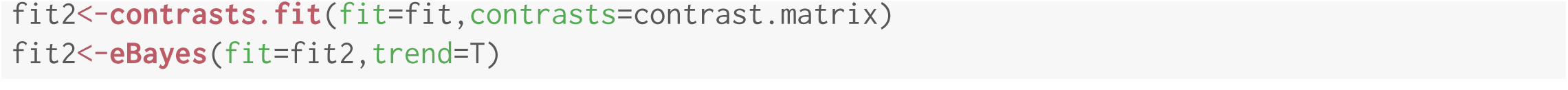

Top ten significant differences in EnzPos vs EnzNeg:

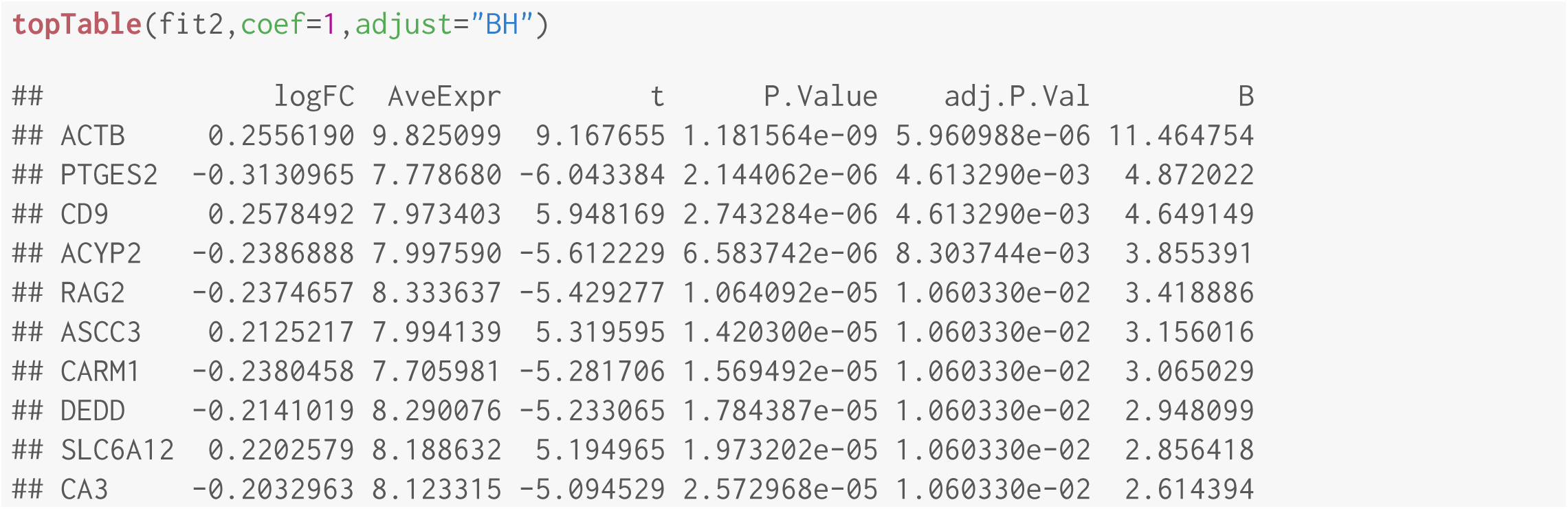

A positive value in the logFC column indicates more counts in Enz-Pos compared to Enz-Neg, and a negative value indicates more counts in Enz-Neg compared to Enz-Pos.

Top ten significant differences in EnzPos vs Initial:

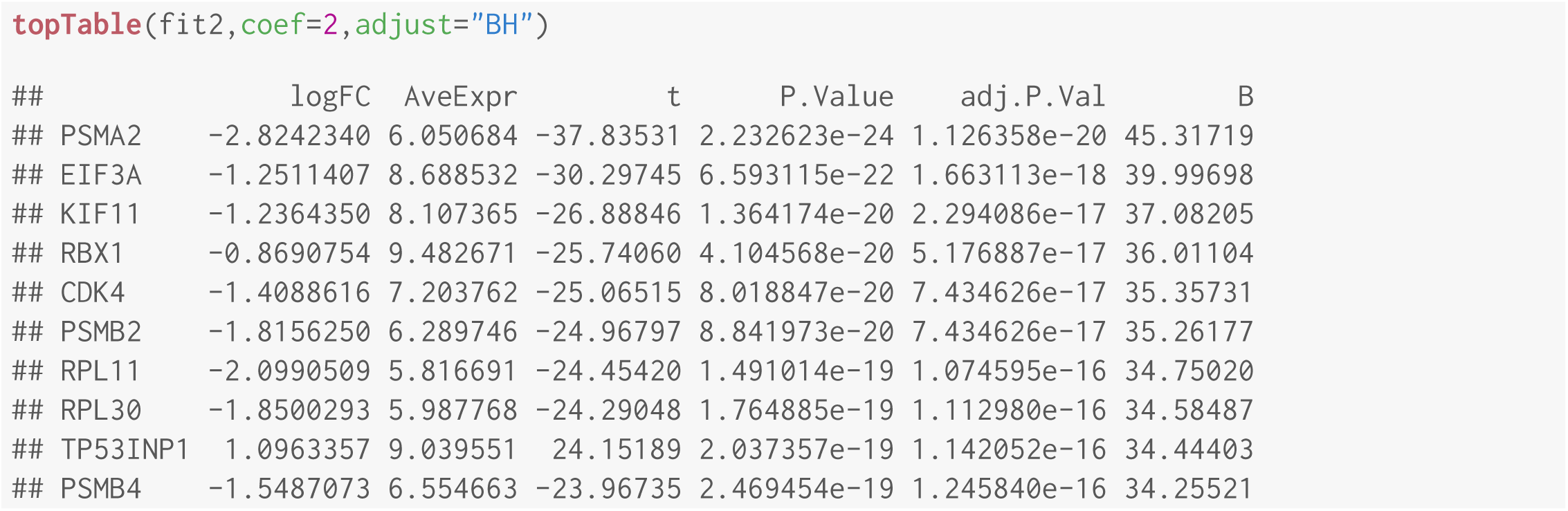

A positive value in the logFC column indicates more counts in Enz-Pos compared to Initial, and a negative value indicates more counts in Initial compared to Enz-Pos.

A given shRNA was considered a “hit” if it showed at least 2-fold abundance decrease in Enz-Pos relative to both Enz-Neg control and initial samples. There are no genes in the first contrast (EnzPos vs. EnzNeg) with a 2-fold abundance decrease, so we tested for any genes with conserved abundance decreases in both sets:

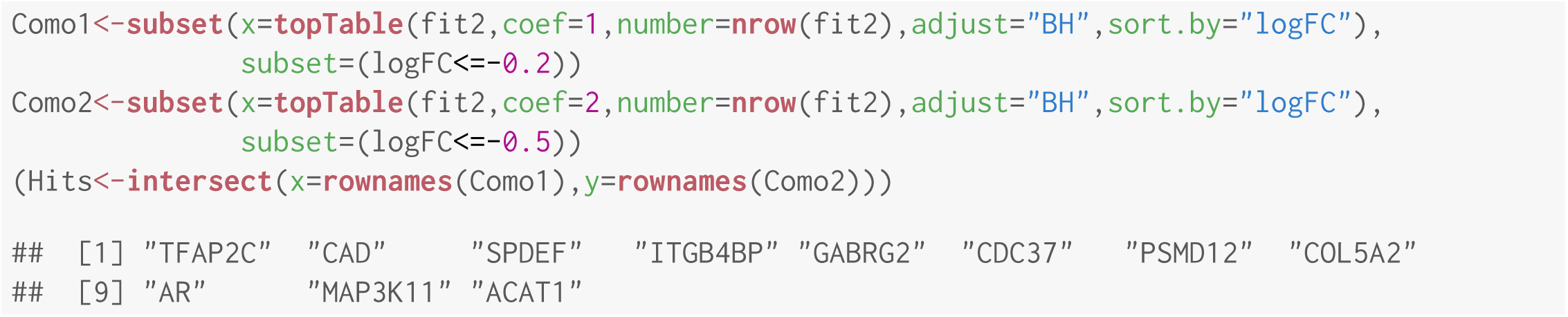

We found 11 genes out of 5045 that satisfy the criteria above to be considered “hits”. The table below shows the log-scaled shRNA counts for the hits:

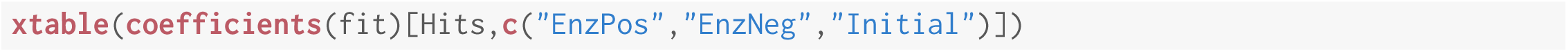

|  | EnzPos | EnzNeg | Initial |
| --- | --- | --- | --- |
| TFAP2C | 6.19 | 6.49 | 7.12 |
| CAD | 5.48 | 5.75 | 6.90 |
| SPDEF | 6.07 | 6.32 | 7.03 |
| ITGB4BP | 6.13 | 6.36 | 7.27 |
| GABRG2 | 7.49 | 7.72 | 8.02 |
| CDC37 | 6.35 | 6.57 | 7.06 |
| PSMD12 | 6.65 | 6.86 | 7.68 |
| COL5A2 | 5.64 | 5.85 | 6.60 |
| AR | 6.79 | 7.00 | 7.70 |
| MAP3K11 | 6.97 | 7.18 | 7.72 |
| ACAT1 | 6.67 | 6.88 | 7.24 |

Here we create two volcano plots, which are convenient diagrams that show the fold changes versus a measure of statistical significance of the change. This code was modified from the native Limma `volcanoplot` code to highlight only our hits.

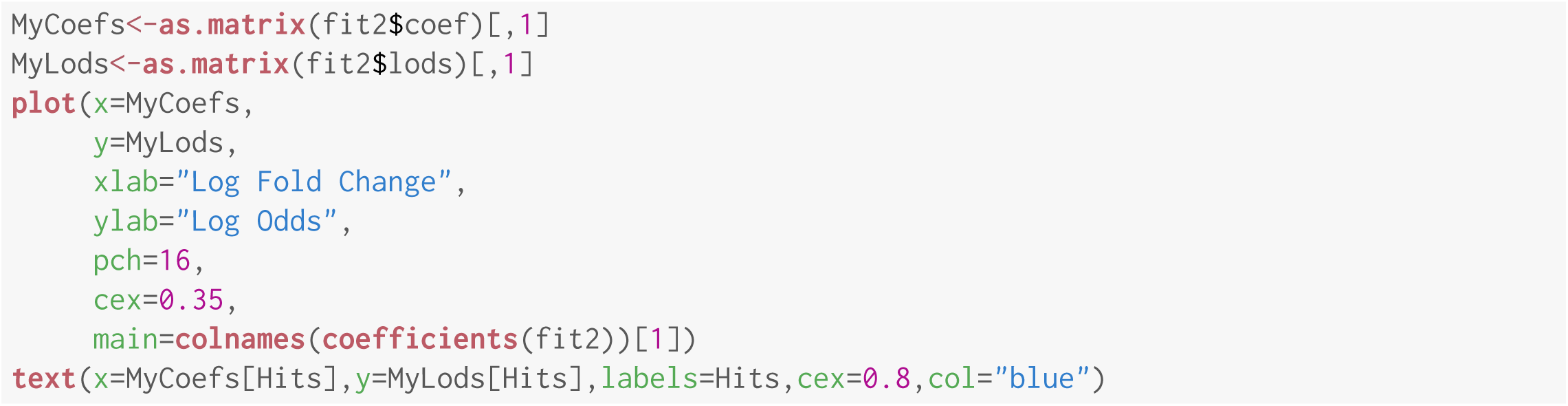

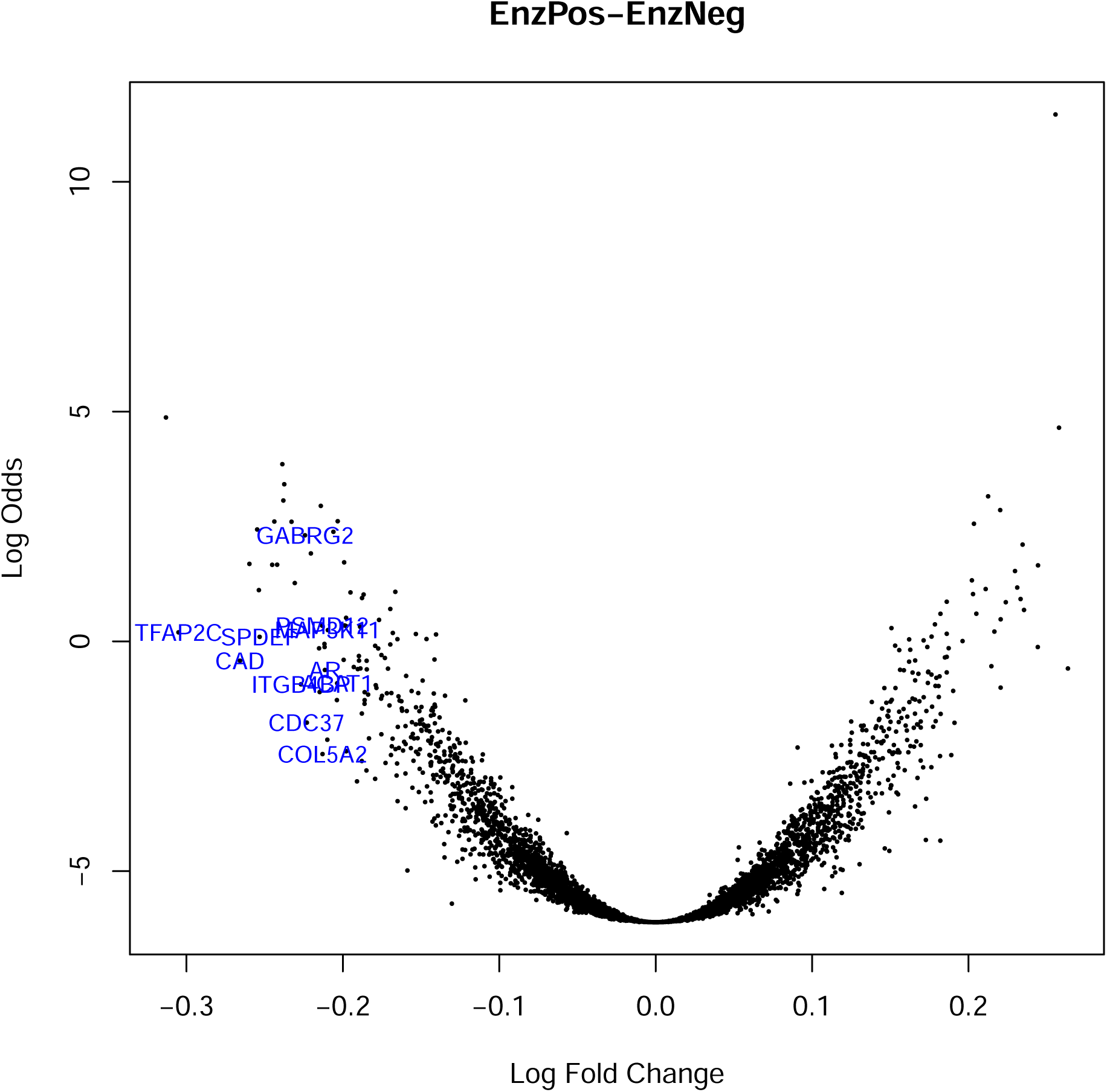

#### Second volcano plot

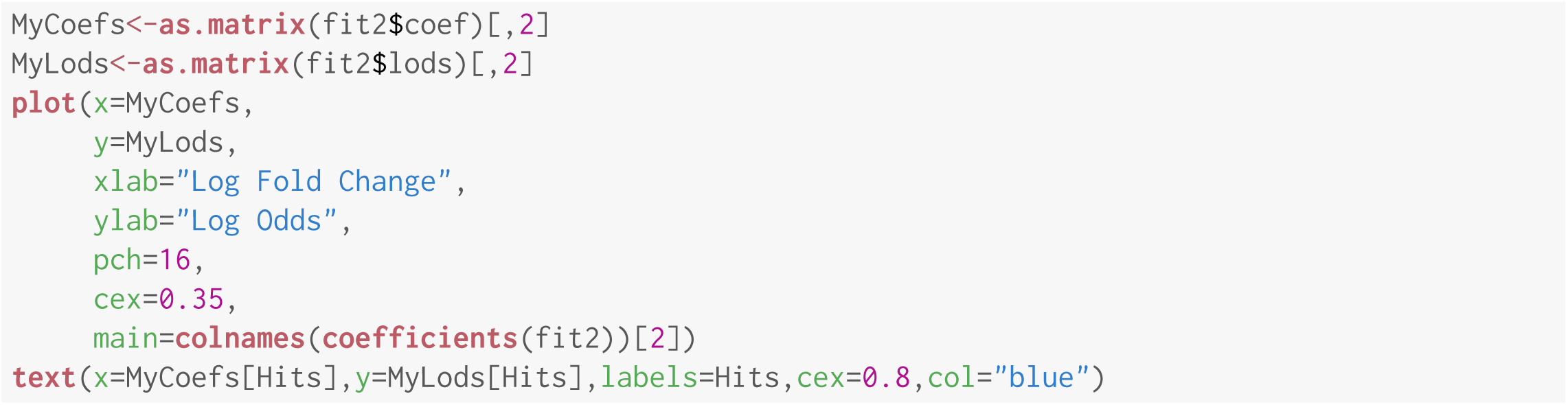

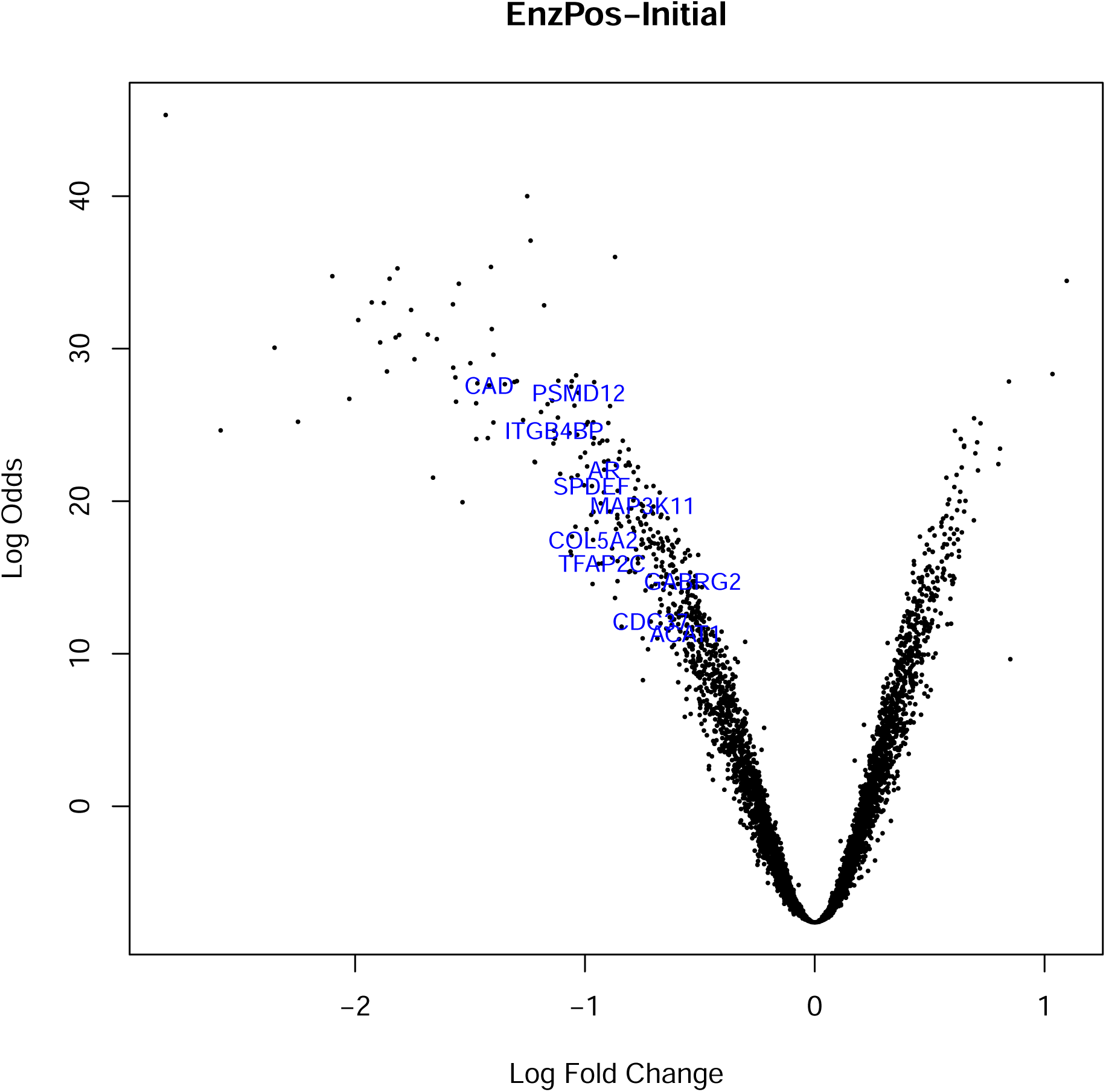

The shRNA count differences are more striking in the EnzPos *vs*. Initial contrast compared to the Enz-Pos *vs*. EnzNeg contrast.

### 5.2 Differential Counts with edgeR’s Negative Binomial Method

We also calculated counts differences using the negative binomial method implemented in edgeR, and achieved identical results as the above test with Limma’s Limma-Trend method.

First we estimated the dispersions:

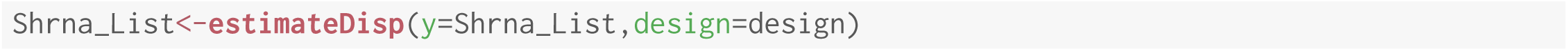

Then we discovered genes with differential counts:

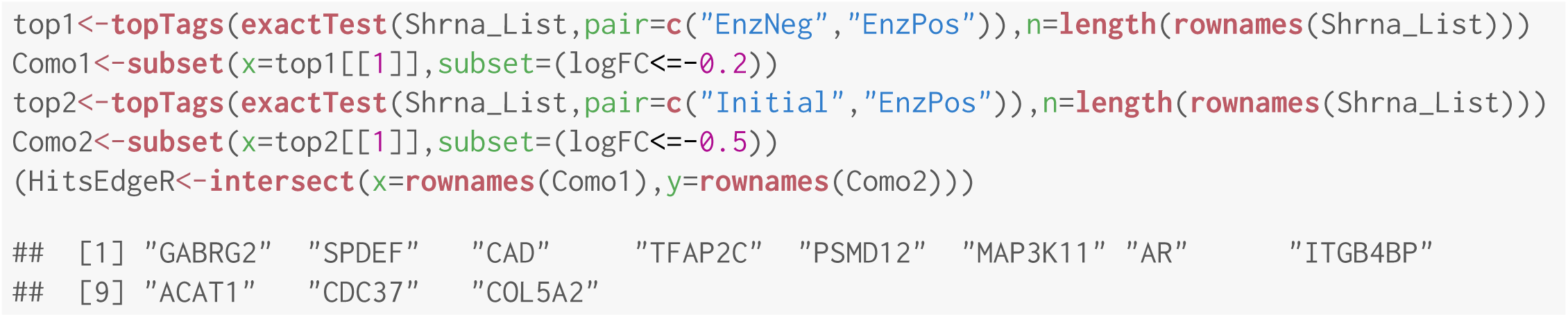

## 6 Compiling Instructions

This document was created using knitR [4] and LATEX 2*ε* in the RStudio Server (http://www.rstudio.com/) integrated development environment (version 0.99.903). Every time the LATEX code is compiled, the code that supports our conclusions is rerun, and in most cases, the results are embedded in this document, facilitating reproducible research. The compile time for this document and its embedded code is approximately nine minutes on our compute server. The .tex and .aux files generated by knitR were compiled to.pdf with `pdflatex` and BibTEX.

### 6.1 Build Citation Database

Here we collect the citation information for all packages and databases we used in this report:

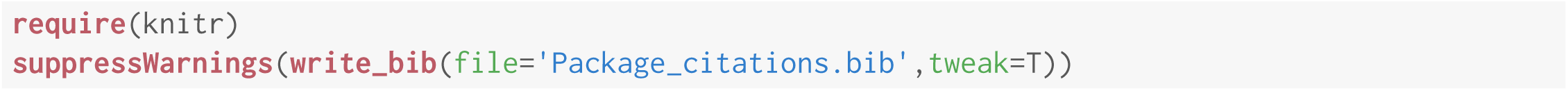

### 6.2 System and Session Information

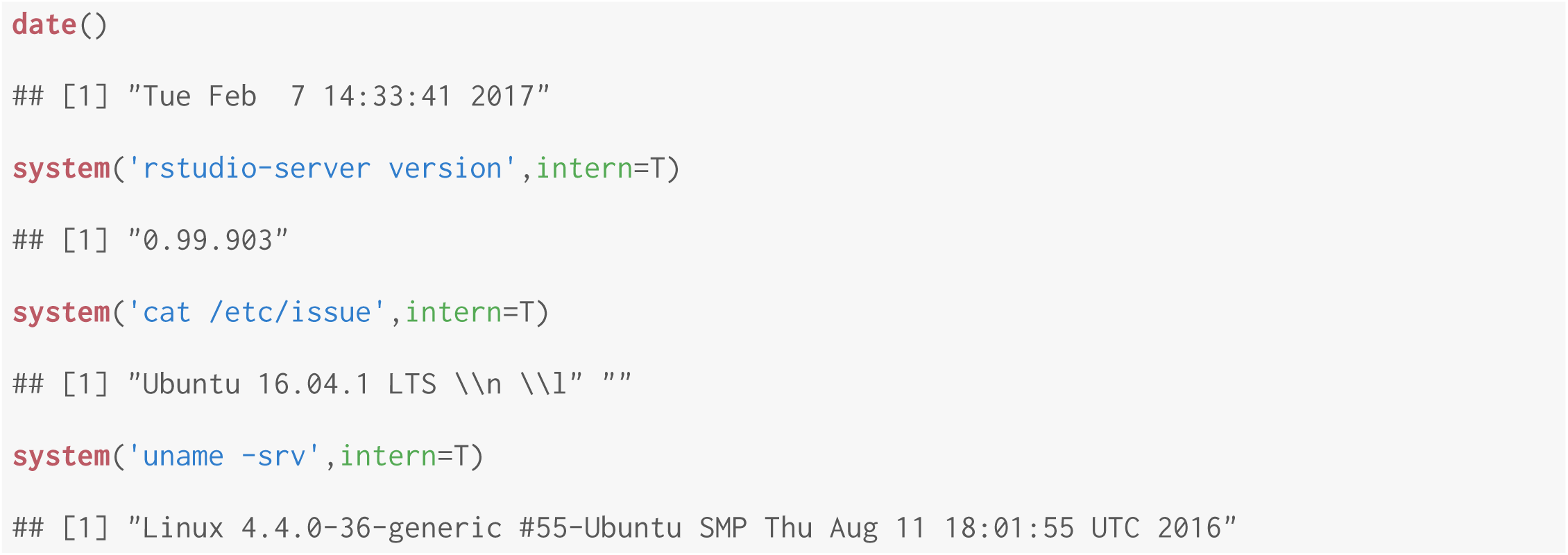

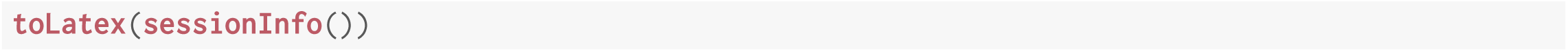

- R version 3.3.1 (2016-06-21), `x86_64-pc-linux-gnu`
- Locale: `LC_CTYPE=en_US.UTF-8`, `LC_NUMERIC=C`, `LC_TIME=en_US.UTF-8`, `LC_COLLATE=en_US.UTF-8`, `LC_MONETARY=en_US.UTF-8`, `LC_MESSAGES=en_US.UTF-8`, `LC_PAPER=en_US.UTF-8`, `LC_NAME=C`, `LC_ADDRESS=C`, `LC_TELEPHONE=C`, `LC_MEASUREMENT=en_US.UTF-8`, `LC_IDENTIFICATION=C`
- Base packages: base, datasets, graphics, grDevices, methods, stats, utils
- Other packages: edgeR 3.14.0, highlight 0.4.7, knitr 1.15.1, limma 3.28.21, plyr 1.8.4, tidyr 0.6.0, xtable 1.8-2
- Loaded via a namespace (and not attached): assertthat 0.1, evaluate 0.10, grid 3.3.1, highr 0.6, lattice 0.20-34, locfit 1.5-9.1, magrittr 1.5, Rcpp 0.12.7, splines 3.3.1, stringi 1.1.2, stringr 1.1.0, tibble 1.2, tools 3.3.1

